# The endo-lysosomal system drives lumen formation in a human epiblast model

**DOI:** 10.1101/2025.08.04.668503

**Authors:** Anusha Rengarajan, Sicong Wang, Chien-Wei Lin, Amber E. Carleton, Nikola Sekulovski, Linnea E. Taniguchi, Mara C. Duncan, Kenichiro Taniguchi

**Affiliations:** Department of Cell Biology, Neurobiology and Anatomy, Medical College of Wisconsin, Milwaukee, WI, USA; Division of Biostatistics, Data Science Institute, Medical College of Wisconsin, Milwaukee, WI, USA; Department of Cell and Developmental Biology, University of Michigan Medical School, Ann Arbor, Michigan, USA; Department of Pediatrics, Medical College of Wisconsin, Milwaukee, WI, USA; Cancer Center, Medical College of Wisconsin, Milwaukee, WI, USA

## Abstract

The formation of a central lumen in the epiblast is a critical step that occurs during implantation in the human embryo. Lumen formation is accompanied by highly dynamic and complex cargo trafficking in the endo-lysosomal system. However, our understanding of key players and machineries that control this critical trafficking process remains incomplete in the context of epiblast development. Here, we explored endo-lysosomal dynamics that are associated with the generation of the apicosome, the earliest stage of lumen formation in a model of human epiblast development based on human pluripotent stem cells. We uncovered a hybrid early/late endosome compartment as well as a previously unrecognized dynamics of late endosome and lysosome compartments in trafficking podocalyxin (PODXL), a sialomucin glycoprotein that helps to establish and maintain the open lumen, during apicosome formation. To gain molecular insight into these unique hybrid endosome and late endosome/lysosome machineries in PODXL traffic, we used APEX2-based spatial proteomics to identify PODXL-proximity partners during apicosome formation, and identified RAB35, a Rab small GTPase known to control PODXL traffic as well as early and late endosome dynamics, as a key player in controlling apicosome formation. Our results suggest that RAB35 limits excess apicosome formation by promoting the early to late endosome transition as well as lysosome formation, which help to reduce PODXL to a level necessary for single apicosome formation. Overall, this study reveals novel endo-lysosomal mechanisms that contribute to apical membrane morphogenesis in a human model of epiblast formation.

## INTRODUCTION

A key step in early human embryonic development is the formation of the epiblast, a radially organized cyst with an apical central lumen and basal exterior (1–3). This polarized epiblast cyst is essential for subsequent critical developmental events such as amniogenesis and gastrulation (4–6). Previous studies using three-dimensional (3D) human pluripotent stem cell (hPSC)-derived model of human epiblast formation have shown that the first sign of lumen formation in this hPSC-epiblast model is the formation of an intracellular structure called an apicosome ((1, 7), **Fig. 1A**). In an aggregate of cells, apicosomes from multiple cells fuse to initiate lumen formation in the center of the aggregate. Each individual apicosome has characteristics of the apical lumen; its membrane contains microvilli and a primary cilium, and its lumen has high calcium concentration (7). Apicosomes are seen in mouse embryos at a similar stage (7, 8), suggesting that apicosomes may contribute to epiblast formation *in vivo*.

**Figure 1.**
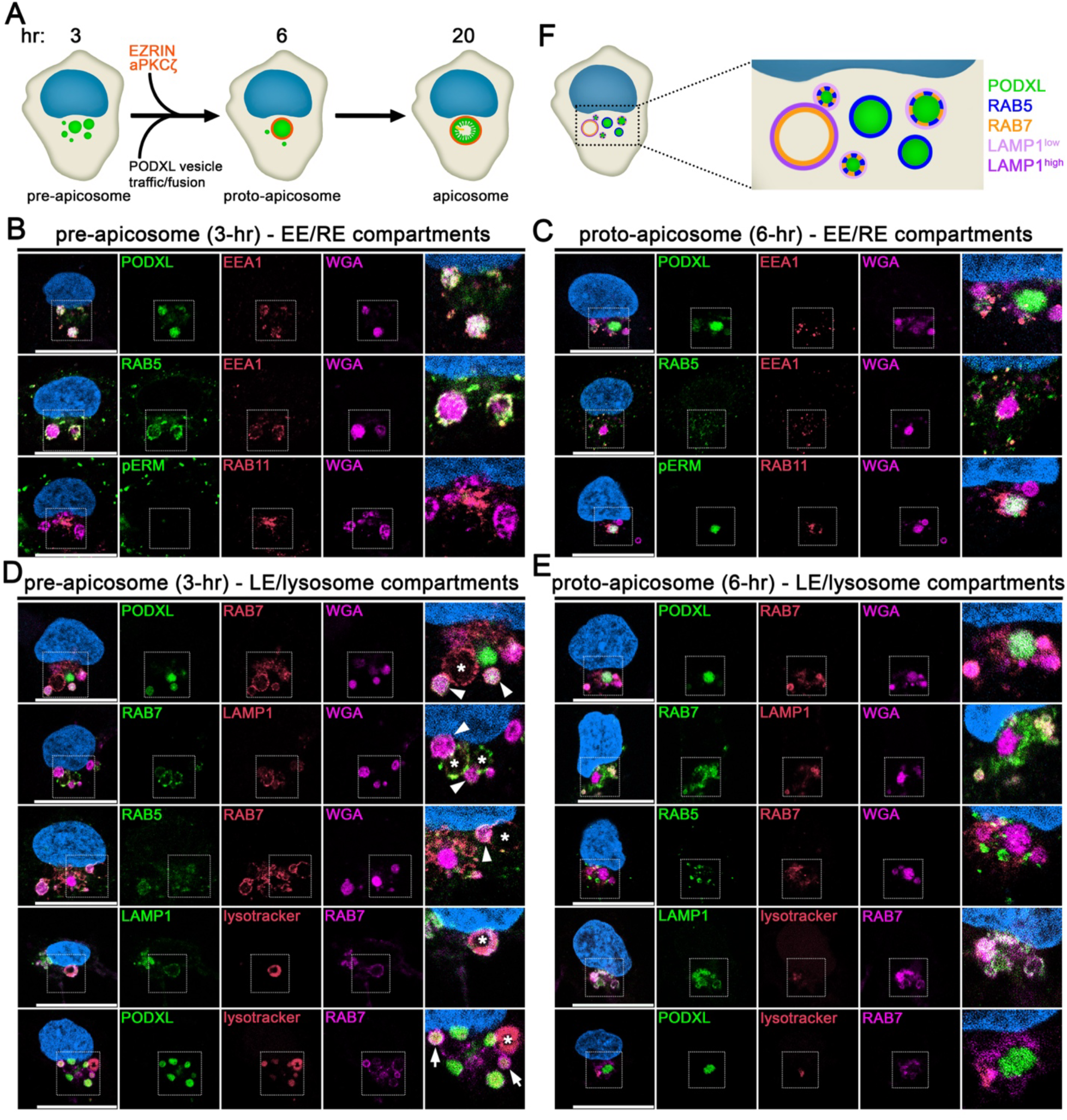
Early endosome and LE/lysosome compartments transiently enlarge during apicosome formation. A) A schematic overview of apicosome formation in hPSC and classification of stages. Pre-apicosome stage occurs first as multiple PODXL vesicles (green) accumulates peri-nuclearly. PODXL vesicle traffic and fusion lead to the recruitment of EZRIN and aPKCζ (red circle) and proto-apicosome formation. Mature apicosome is formed as the proto-apicosome grows in size and acquire additional apical characteristics (e.g., microvilli (green projections), primary cilium (yellow projection), high Ca^2+^ concentration (not shown)). B-E) Representative confocal images of H9 hESC harvested at 3-(B,D) and 6-(C,E) hr timepoints, which were stained with indicated markers (B and C, EE and RE components; D and E, LE/lysosome components). Two additional representative cells are shown in Fig. S2A-D (EE/RE) and S3A-D (LE/lysosome). Arrowheads indicate PODXL/WGA structures that contain RAB7; arrows indicate structures that contain PODXL, RAB7 and lysotracker; asterisks indicate highly enlarged LE/lysosome compartments labeled by RAB7, LAMP1 and/or lysotracker, which lack PODXL/WGA. F) A schematic summarizing molecular characteristics of the endo-lysosomal system during the earliest stages of apicosome formation. Scale = 20µm. Blue pseudocolor indicates DNA in all images.

The process of apicosome formation takes place in three distinct stages (**Fig. 1A**). First, vesicles containing apical proteins such as podocalyxin (PODXL), a member of the CD34 sialoprotein family that is a critical driver of lumenogenesis in several systems (9–14), accumulate peri-nuclearly (pre-apicosome stage, first 3 hours after initiation). Then, actin cytoskeleton and additional apical proteins (e.g., EZRIN, aPKCζ) accumulate to form nascent apicosomes (proto-apicosome stage, between 6 and 12 hours after initiation). Finally, the proto-apicosome matures through an increase in volume and the formation of microvilli and a primary cilium (mature apicosome stage, between 12 and 20 hours after initiation (1, 7), **Fig. 1A**).

We previously reported that the apicosome forms near markers of the early, recycling and late endosomes, as well as the lysosome (1, 7, 15). Moreover, the forming apicosome transiently associates with endocytosed material (7), suggesting that the endo-lysosomal system may contribute to apicosome formation. However, the exact timing and functional significance of the endo-lysosomal pathway in apicosome formation remained unclear. Here, we explored endo-lysosomal dynamics during apicosome formation using immunofluorescence (IF) and spatial proteomics using the engineered ascorbate peroxidase APEX2 (16, 17). We uncovered a series of previously unrecognized trafficking events, driven by a RAB5/RAB7 hybrid endosome, that occur at the pre-apicosome stage to ensure that only one apicosome is formed per cell. Overall, these results present a novel model of apical morphogenesis, which depends on the endo-lysosomal pathway that is used during early human development.

## RESULTS

### The endo-lysosomal system undergoes dynamic remodeling during apicosome formation

To gain insight into the roles of the endo-lysosomal system during apicosome formation, we first monitored the localization of endo-lysosomal organelles at different apicosome formation stages (pre-apicosome, 3-hour (hr); proto-apicosome, 6-hr; mature apicosome, 20-hr, see **Fig. S1A,B** for IF images and quantitation for different apicosome stages at several timepoints) in control cells (H9 human embryonic stem cell line) by using IF. Importantly, apicosome formation is asynchronous (**Fig S1B**), because the cells are still undergoing mitosis. Therefore, we identified apicosome stages based on the size, morphology and number of PODXL vesicles and/or the presence or absence of pERM (phosphorylated EZRIN/RADIXIN/MOESIN) as described in Materials and Methods. Wheat germ agglutinin (WGA) was also used to label PODXL vesicles. WGA binds to glycosylated proteins like PODXL (18–20), and co-localizes with PODXL under all conditions in this study (**Fig. 1B-E**).

We first monitored the early endosome (EE) using antibodies against EEA1 (early endosome antigen 1) and RAB5. In the pre-apicosome stage, cells contained two to three enlarged (>1μm at the widest diameter) EEs. These enlarged EEs were almost always positive for PODXL/WGA (**Fig. 1B**, see **Fig. S2A,B** for additional sets of images). These EEs were also in close proximity to the nucleus (**Fig. 1B**), consistent with the known association of the apicosome with the centrosome (7). In addition to the enlarged EE, cells in the pre-apicosome stage contained several small peripheral EEs (**Fig. 1B**). These peripheral EEs were also often labeled by PODXL/WGA (**Fig. 1B**), suggesting that PODXL and other apicosome-directed proteins transit through the EE at this stage.

In contrast to the enlarged EEs in the pre-apicosome stage, enlarged EEs were absent in the proto-apicosome stage (**Fig. 1C**, see **Fig. S2C,D** for additional sets of images). Instead, several small EEs were observed adjacent to the proto-apicosome (**Fig. 1C**). Notably, PODXL and WGA were absent from the EEs at this stage, regardless of whether the EEs were adjacent to the proto-apicosome or more peripheral (**Fig. 1C**). Finally, at the apicosome stage, EEs were seen adjacent to the apicosome, and were also seen in the cell periphery (**Fig. S2E,F**). Similar to the proto-apicosome stage, PODXL and WGA were not found in the EE at the apicosome stage (**Fig. S2E,F**). Together, these results suggest that, during the early stages of apicosome formation, apicosomal material transits through the EE. Furthermore, this transit is associated with a transient enlargement of the early endosomes, suggesting that the early endosome itself may contribute to the content of the apicosome.

We next examined the association of the recycling endosome (RE) with the forming apicosome. We found that RE labeled with RAB11 was concentrated near the nucleus proximal to the enlarged PODXL/WGA positive compartments at both the pre- and proto-apicosome stages (**Fig. 1B,C**, also see **Fig. S2A-D**). At the apicosome stage, the RE was closely associated with one side of the apicosome, likely near the base of the primary cilium (**Fig. S2E,F**). These findings suggest that the location or morphology of the RE is not altered during apicosome formation.

We next explored the late endosome (LE) and lysosome. Because these compartments share many characteristics (21–23), and, therefore, are often difficult to distinguish, especially in dynamic contexts, the presence of the small GTPase RAB7, the membrane protein LAMP, and/or lysotracker, a dye that stains acidic compartments was used to identify compartments that could either be the LE or lysosome (LE/lysosome). At the pre-apicosome stage, several enlarged LE/lysosomes containing PODXL/WGA were observed (**Fig. 1D**, see arrows and arrowheads, see **Fig. S3A,B** for additional images). Notably, most cells also contained one or more highly enlarged structures (>2μm, widest diameter) that were labeled with RAB7, LAMP1 and lysotracker, but were devoid of PODXL/WGA (**Fig. 1D**, asterisks). Finally, unlike the EE, small peripheral LE/lysosome compartments were rarely observed in these cells.

In the proto-apicosome stage, cells contained enlarged LE/lysosome compartments that were positive for PODXL, albeit much fewer than in the pre-apicosome stage; however, the highly enlarged LE/lysosomes that lack PODXL staining were absent (**Fig. 1E**, see **Fig. S3C,D** for additional images). Notably, at the proto-apicosome stage, small LE/lysosomes accumulated adjacent to the enlarged proto-apicosome labeled with PODXL and WGA (**Fig. 1E**). Finally, at the apicosome stage, cells lacked enlarged LE/lysosomes (**Fig. S3E,F**). These results indicate that, like the EEs, the LE/lysosomes undergo a transient enlargement. Moreover, the enlarged LE/lysosomes persist longer than the enlarged EEs.

Finally, we examined the relationship between the EE and LE/lysosome during apicosome formation. In the pre-apicosome stage, all PODXL/WGA structures contained markers of the EE, while only a subset of PODXL/WGA structures contained markers of the LE/lysosome (**Fig. 1B**). Accordingly, EE and LE/lysosome components co-localized on a subset of enlarged PODXL/WGA structures (**Fig. 1B,D**), suggesting that these enlarged PODXL/WGA structures are hybrid organelles with characteristics of both the early and late endo-lysosomal system (**Fig. 1F**). In contrast, at the proto-apicosome stage, although EE, LE and lysosomal components are seen adjacent to the proto-apicosome, these proteins no longer co-localized (**Fig. 1B-E**).

Moreover, the remaining enlarged LE and lysosome compartments lacked RAB5 (**Fig. 1E**). Together, these results reveal a complex transient reorganization of the endo-lysosomal system that accompanies apicosome formation.

### The PODXL-proximity proteome reveals machinery important for endo-lysosomal dynamics

To gain additional insight into machinery controlling endo-lysosomal dynamics during apicosome formation, we examined the proteome of the PODXL^+^ endosomes and apicosomes at 30-hr, a timepoint when all three stages of apicosome maturation can be observed (**Fig. S1B**). To do this, we used APEX2-based proximity labeling using doxycycline (DOX)-inducible PODXL constructs C-terminally tagged with APEX2 (PODXL-APEX2), and APEX2 fused to a nuclear export sequence (APEX2-NES) for subsequent background subtraction during ratiometric analysis (**Fig. 2A,B**, (15, 24)).

**Figure 2.**
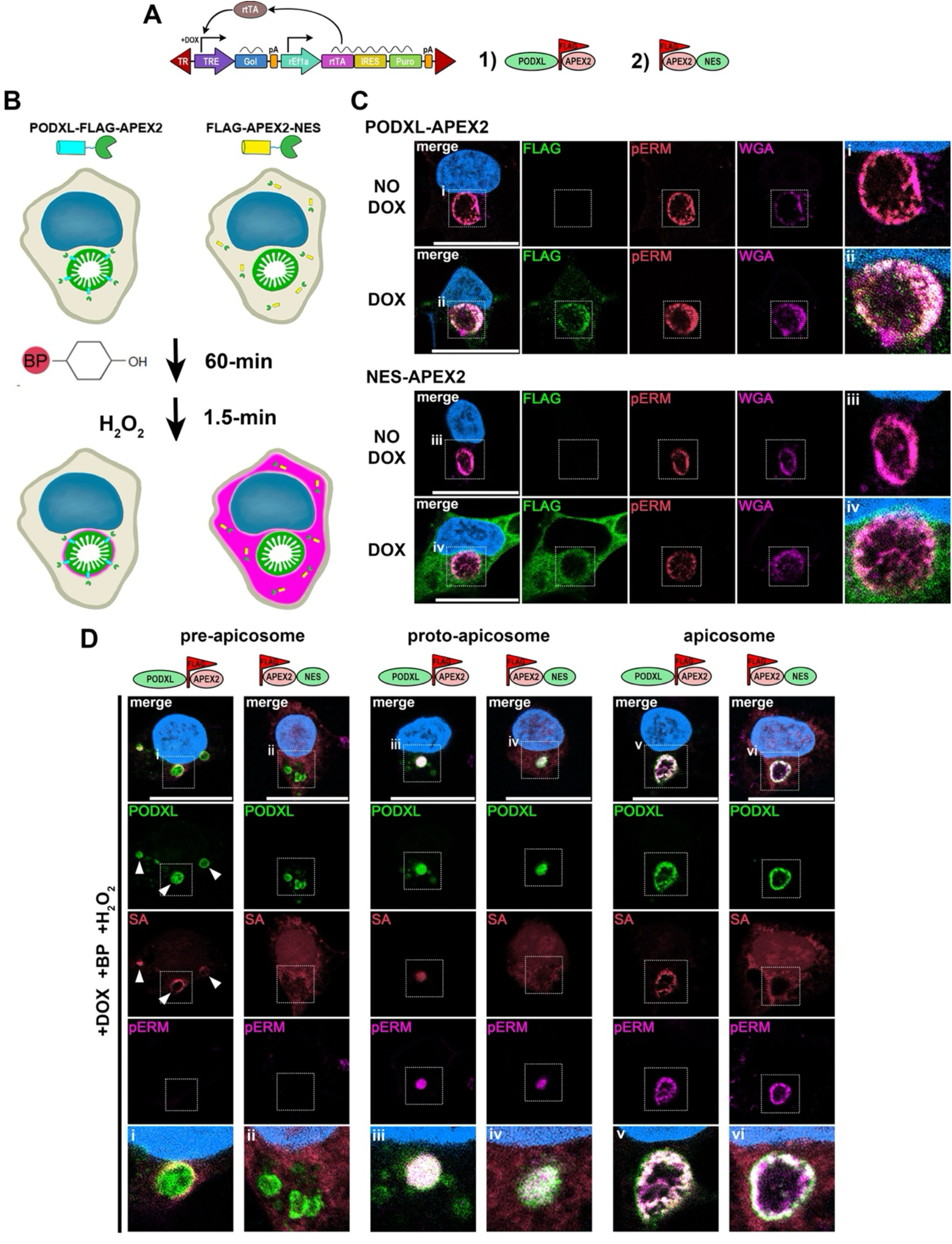
APEX2-based spatial biotinylation of the PODXL proximal territory during apicosome formation. A) Design of APEX2 fusion construct based on a DOX-inducible piggyBac transposon system. GoI, gene of interest; TR, terminal repeats; TRE, tetracycline-responsive element; pA, polyadenylation sequence; rEf1α, rat elongation factor 1 α promoter; rtTA, reverse tetracycline transactivator; IRES, internal ribosome entry site; Puro, puromycin. Resulting proteins are shown on the right (1) PODXL-FLAG-APEX2; 2) FLAG-APEX2-NES). B) Flow diagram of APEX2-based proximity biotinylation of PODXL and cytosolic territory proteins. H9 hESC expressing APEX2 fused to PODXL and NES were incubated with BP (biotin-phenol) for 1 hour, followed by brief H_2_O_2_ treatment to trigger biotinylation. Purple shaded area indicates biotinylated territory. C) Representative optical section of 20-hr cells expressing PODXL-or NES-APEX2 constructs with or without DOX, stained with indicated markers. With DOX treatment, FLAG signal is seen in the pERM^+^ apicosome in the PODXL-APEX2 cells, while abundant cytosolic FLAG staining is seen in the NES-APEX2 cells. D) Representative confocal images of biotin-labeled PODXL-or NES-APEX2 cells from pre-apicosome, proto-apicosome and mature apicosome stages stained with indicated markers. Arrowheads indicate streptavidin-labeled PODXL vesicles. In the fluorescent images, insets ((i)-(iv) in C; (i)-(vi) in D) indicate magnified regions in merge. Scale = 20µm. Blue pseudocolor indicates DNA in all images.

We established and validated stable cell lines with each APEX2 construct. To validate that the APEX2 constructs are localized correctly, we induced transgene expression by adding DOX (2µg/mL) to the cells when they were plated (at 0-hr). The expression of PODXL-APEX2 and APEX2-NES was detected by IF. PODXL-APEX2 was enriched at the apicosome territory whereas APEX2-NES was observed throughout the cytoplasm based on FLAG staining (**Fig. 2C**). We next performed APEX2 labeling reactions and monitored the enrichment of biotinylated species with fluorophore-conjugated streptavidin, which binds to biotin. APEX2 labeling was performed by incubating cells with biotin-phenol (BP) for 1 hour, followed by 1.5 minute H_2_O_2_ labeling and harvesting (**Fig. 2B**). In control samples (with no BP or H_2_O_2_), weak diffuse staining was observed (**Fig. S4**). In contrast, in cells expressing PODXL-APEX2, strong biotin signal was detected in the peri-nuclear vesicles (pre-apicosome, **Fig. 2D**, arrowheads) and at the proto- and mature apicosomes (**Fig. 2D**), whereas, in cells expressing APEX2-NES, strong biotin signal was detected in the entire cytoplasmic domain (**Fig. 2D**). For PODXL-APEX2, although labeling was concentrated at the apicosome, additional diffuse cytosolic localization of biotin signal was observed, likely due to diffusion of the peroxidase activated label, or diffusion of labeled cytosolic proteins (**Fig. 2D**). These analyses demonstrate that this selective APEX2-based biotinylation can label and enrich PODXL-proximity proteins in apicosome-forming cells.

To identify the PODXL-proximity proteome, we isolated biotinylated proteins from PODXL-APEX2 and NES-APEX2 cells in duplicate, a non-biotinylated APEX2 negative control (cells expressing PODXL-APEX2 without H_2_O_2_ labeling), and a no APEX2 negative control (unmodified H9 hESC with H_2_O_2_ labeling, **Fig. 3A,B**). Isolated proteins were subjected to on-bead digestion followed by tandem mass tag (TMT)-based tagging. The TMT-tagging allows a ratiometric analysis of samples to improve the specificity of protein identification from complex samples (25, 26).

**Figure 3.**
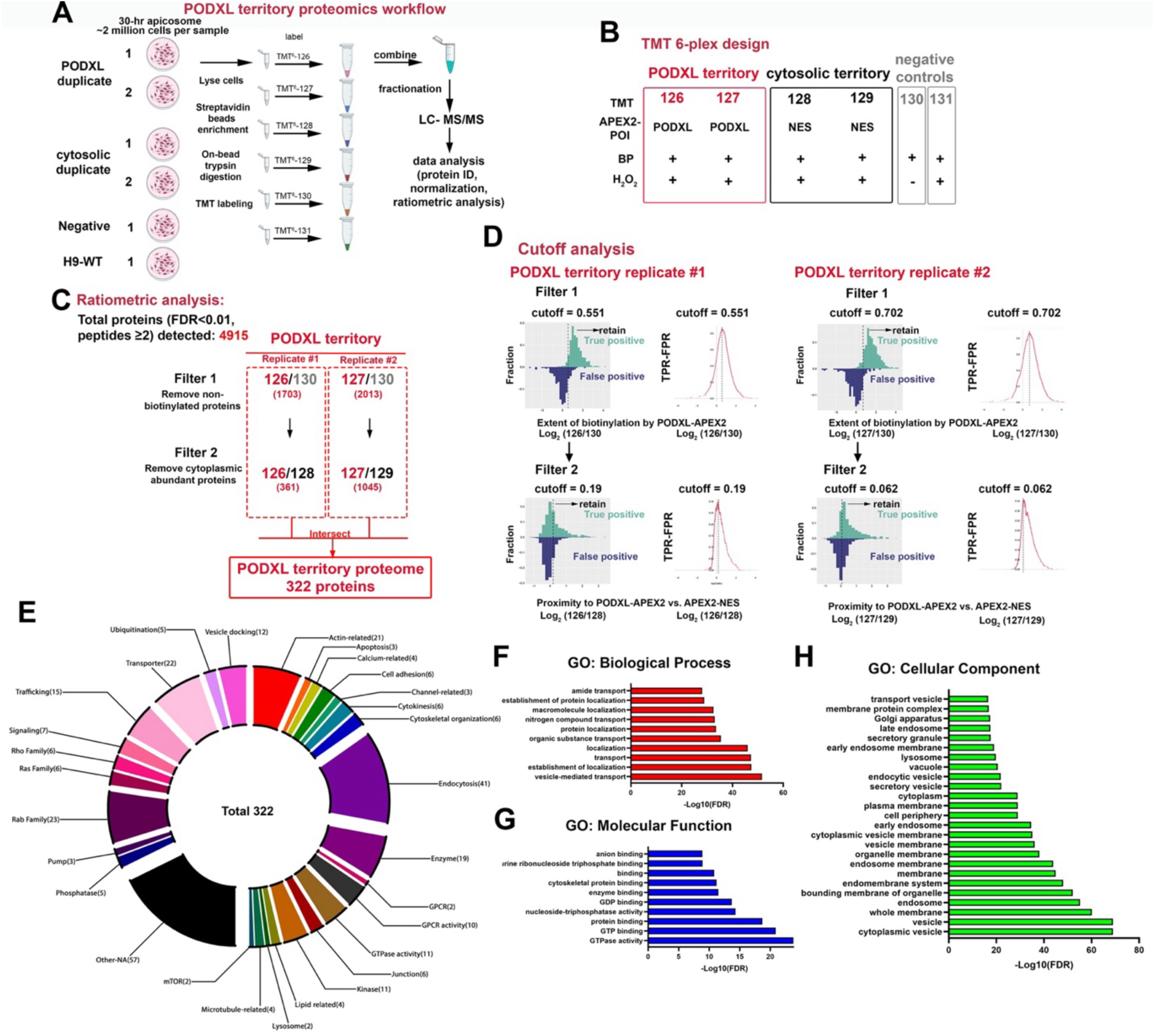
Ratiometric profiling of PODXL-proximity proteome during apicosome formation. A) Workflow of PODXL territory proteome analysis using duplicates of PODXL and NES samples as well as two separate negative controls (PODXL-APEX2 without H_2_O_2_, unmodified H9 treated with BP and H_2_O_2_). B) Experimental design of TMT 6-plex-based profiling. Row 1, specific TMT tags; row 2, APEX2 lines; rows 3 and 4, proximity labeling conditions. POI, protein of interest. C) Diagram of ratiometric analyses to obtain proteins specific to PODXL territory. Brackets: Number of proteins that passed each filter. D) Representative histograms and receiver operating characteristic (ROC) curves of PODXL replicate 1 and 2, illustrating how filters 1 and 2 were applied. See Materials and Methods for additional cutoff analysis details. E) Functional classification of the newly identified PODXL territory proteins. F-H) Gene Ontology analysis of the PODXL territory proteins: Biological Process (F, top 10 shown), Molecular Function (G, top 10) and Cellular Component (H, top 25).

All samples were then combined, fractionated, and analyzed as a pooled mixture by LC-MS/MS (**Fig. 3A**). We identified 4,915 unique proteins (**Fig. 3C**, see **Table 1A** for the raw dataset). Using our previously established ratiometric analysis pipeline ((15), **Fig. 3C,D**, also described in detail in Materials and Methods), this dataset was normalized based on the enrichment of known mitochondrial matrix soluble proteins (**Table 1B**, (16, 17, 24)). To identify proteins selectively enriched in the PODXL-proximity territory, the dataset was filtered based on the level of known proteins with cell membrane annotations in uniport (**Table 1C**) in the PODXL-APEX, NES-APEX and negative control samples (**Fig. 3C,D**). Followed by filtering (protein list after each filtering is shown in **Table 1D-G**), this analysis identified 322 proteins (**Table 2A**) and accurately identified 14 out of 82 known PODXL binding partners based on BioGrid (**Table 2B**, 51 of those 82 protein coding genes were expressed in hPSC, (27)), suggesting that the identified proteins represent a *bona fide* PODXL-proximal proteome.

**Table 1.** PODXL territory proteome dataset and list of proteins after Filter 1 and Filter 2. A) PODXL-APEX2 6-plex MS dataset. B) List of 495 mitochondrial matrix proteins C) Uniprot human protein database used in this study D-G) List of proteins after Filter 1 (D,E) and Filter 2 (F,G): PODXL-APEX2 #1 list (D,F); PODXL-APEX2 #2 (E,G).

**Table 2.** PODXL territory proteome during apicosome formation. A) List of 322 PODXL proximal proteins after intersecting the post-Filter 2 lists from samples #1 and #2. B) Interaction analysis for the 322 PODXL proximal proteins. Fourteen known PODXL binding partners were identified using BioGrid (out of 82 proteins). Among the 82 proteins, 51 of the 82 protein coding genes were expressed. C) UniProt- and GO-based functional annotation of the 322 PODXL territory proteins. D-F) GO analysis of the 322 PODXL proximal proteins (Biological Process, D; Molecular Function, E; Cellular Component, F).

Our PODXL proximity proteome contained proteins annotated to localize to the EE, RE, LE, consistent with our imaging data (**Fig. 1**, **Table 2A**). Additionally, functional annotation of 322 proteins reveals enrichment in proteins associated with a spectrum of molecular and cellular machineries such as actin cytoskeletal organization, endocytosis and GTPase activity (Rab, Ras, Rho family, **Fig. 3E**, also see **Table 2C**). Indeed, many of these functional machineries (**Fig. 3F,G**, e.g., GTPase activity, vesicle traffic) as well as subcellular localization (**Fig. 3H**, e.g., EE, LE, lysosome) were also identified in the Gene Ontology (GO) enrichment analysis. These data confirm the localization of PODXL in many compartments of the endo-lysosomal system during apicosome formation observed by IF.

### RAB35 regulates apicosome formation

We identified RAB35 among the endo-lysosomal transport proteins enriched in the PODXL-proximal proteome. RAB35 is a known regulator of PODXL trafficking in MDCK cells (13, 28–30). Therefore, we examined its role in apicosome assembly. We first examined its localization in a time course during apicosome formation. In cells at the pre-apicosome stage, all PODXL vesicles contained RAB35 regardless of whether they contained markers of the LE/lysosome (**Fig. 4A**, see arrowheads for RAB7^+^ PODXL vesicles). However, RAB35 was absent from highly enlarged PODXL-negative RAB7^+^/LAMP1^+^ LE/lysosomes (**Fig. 4A**, asterisks). In cells with a proto-apicosome or mature apicosome, RAB35 was localized to the apicosome membrane, as defined by pERM staining (**Fig. 4B,C**). This specific localization to the pERM-positive proto-apicosome structure is distinct from other tested RABs (**Fig. 1**). These results identify RAB35 as an apicosome membrane RAB.

**Figure 4.**
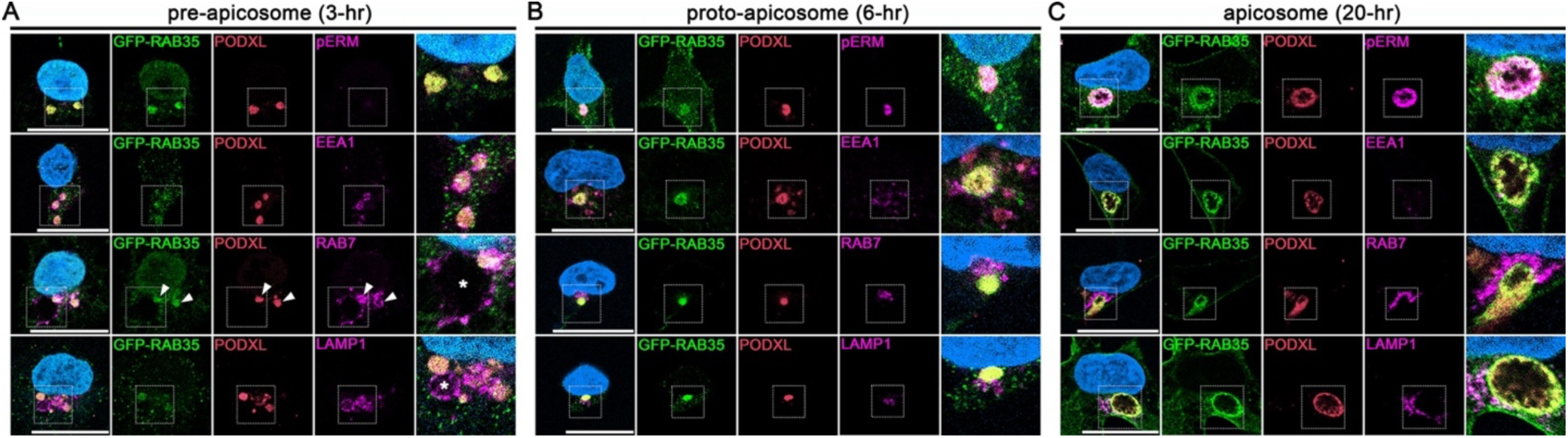
RAB35 co-localizes with apicosomes during formation and after maturation. A-C) H9 hESC carrying DOX-inducible GFP-RAB35-WT transgenic construct were treated with DOX starting at 0-hr, harvested at indicated timepoints (3-hr, (A); 6-hr, (B), 20-hr, (C)), and were stained with indicated markers. Representative confocal images of these samples are shown. Arrowheads indicate RAB7-labeled PODXL vesicles. Asterisks in (A) indicate highly enlarged LE/lysosome compartments that lack PODXL. Insets indicate magnified regions in the merged images. Scale = 20µm. Blue pseudocolor indicates DNA in all images.

Next, to investigate the role of RAB35 in apicosome formation and endosome dynamics, we used two independent hPSC lines lacking RAB35 (one line previously established in (15), and new line established for this study, **Fig. S5**). In both *RAB35*-KO lines, apicosome formation was severely disrupted. At 20-hr, *RAB35*-KO cells failed to form a single apicosome structure (**Fig. 5**). Instead, several apicosome-like structures that were positive for PODXL and pERM were formed (**Fig. 5A**, quantitation shown in **Fig. 5B,S6A**). In addition, in the *RAB35*-KO background, primary cilia formation was impaired. In controls, a well-formed primary cilia (visualized with ARL13B) was present in nearly every mature apicosome (**Fig. 5C**). However, in the *RAB35*-KO cells, although ARL13B staining was observed associated with one of the pERM^+^ structures, the staining was often diffuse or the cilia were notably shorter compared to controls (**Fig. 5C**). Both the multiple apicosome and primary cilia phenotypes were rescued when a wild-type form of RAB35 fused to GFP at its N-terminus (GFP-RAB35) was expressed in the *RAB35*-KO background (**Fig. 5A-C**). These data indicate that RAB35 is critical for apicosome formation and maturation.

**Figure 5.**
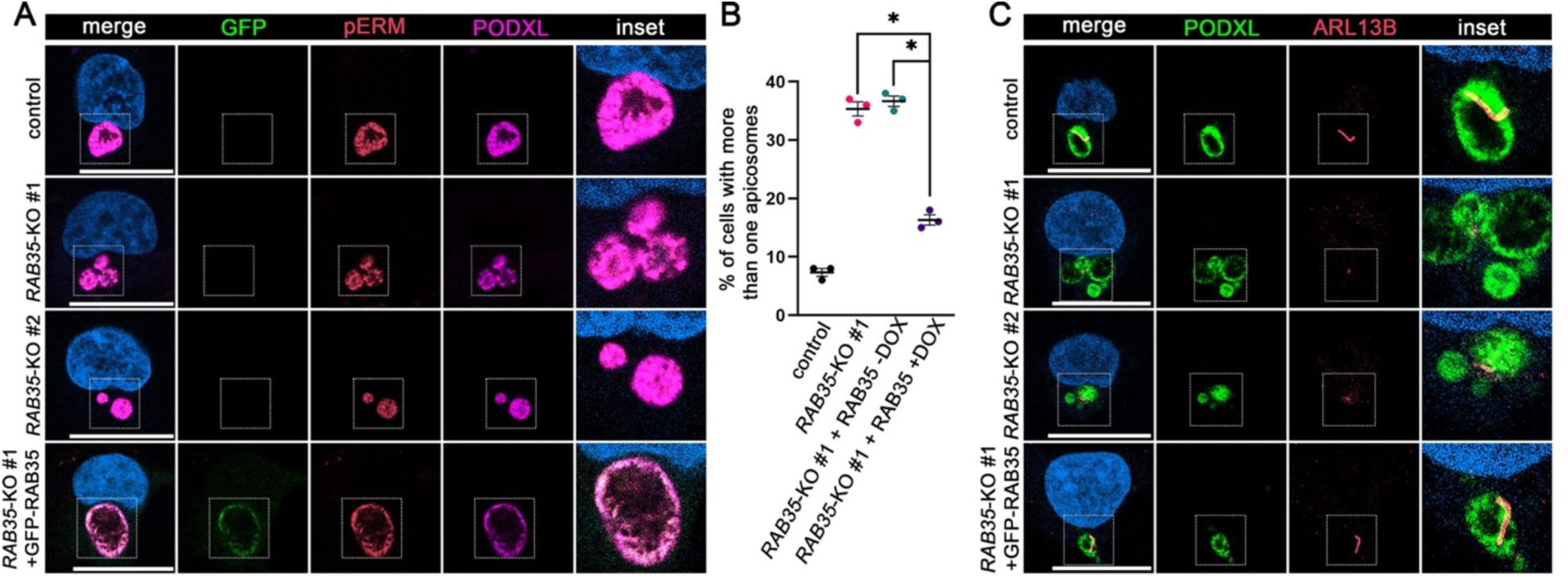
Loss of RAB35 leads to the formation of several apicosomes. A) Representative confocal images of control, *RAB35*-KO #1 and *RAB35*-KO#2 cells as well as *RAB35*-KO cells expressing a DOX-inducible GFP-RAB35-WT construct (*RAB35*-KO + GFP-RAB35) at 20-hr, stained with indicated markers. Formation of several apicosomes (labeled by both PODXL and pERM) was seen in the absence of RAB35, which was rescued when RAB35 was re-introduced. B) Quantitation for the formation of the multi-apicosome phenotype in control (unmodified H9 hESC, black), *RAB35*-KO (red), *RAB35*-KO + GFP-RAB35-WT without (green) or with (purple) DOX treatment. RAB35 re-introduction in the *RAB35*-KO background leads to a significant reduction in multiple apicosome formation (n = 100 cells in each of the three independent experiments in each background, statistical significance based on one-way ANOVA (p < 0.05) as well as by Tuckey’s test (asterisks indicate statistical significance of selected pairwise comparisons, p < 0.05)). C) Loss of RAB35 impairs cilia formation. Representative confocal images of control, *RAB35*-KO #1 and *RAB35*-KO #2 as well as *RAB35*-KO cells expressing a DOX-inducible GFP-RAB35-WT construct (*RAB35*-KO + GFP-RAB35) at 20-hr, stained with indicated markers. Scale = 20µm. Blue pseudocolor indicates DNA in all images.

### RAB35 impairs late endo-lysosomal dynamics at the pre-apicosome stage

We next explored endosome dynamics in *RAB35*-KO cells. At the pre-apicosome stage, components of the EE were associated with enlarged PODXL vesicles similar to control cells (**Fig. 6A**, 3-hr cells are shown). Similarly, LE/lysosome components were associated with PODXL vesicles in the *RAB35*-KO cells (**Fig. 6B**). However, in the *RAB35*-KO cells, the highly enlarged structures that were labeled with LE/lysosome markers but lacked PODXL/WGA were absent (**Fig. 6B**). Instead, *RAB35*-KO cells occasionally displayed a much smaller structure that was labeled with LE/lysosome markers but lacked PODXL/WGA (**Fig. 6B**, asterisk, see **Fig. S6B** for LAMP1^+^ vesicle size). These results suggest that traffic to the LE/lysosome may be impaired in the *RAB35*-KO cells. Consistent with this hypothesis, levels of PODXL were elevated in the *RAB35*-KO cells, whereas levels of RAB7 were unaffected (**Fig. 6C-E**).

**Figure 6.**
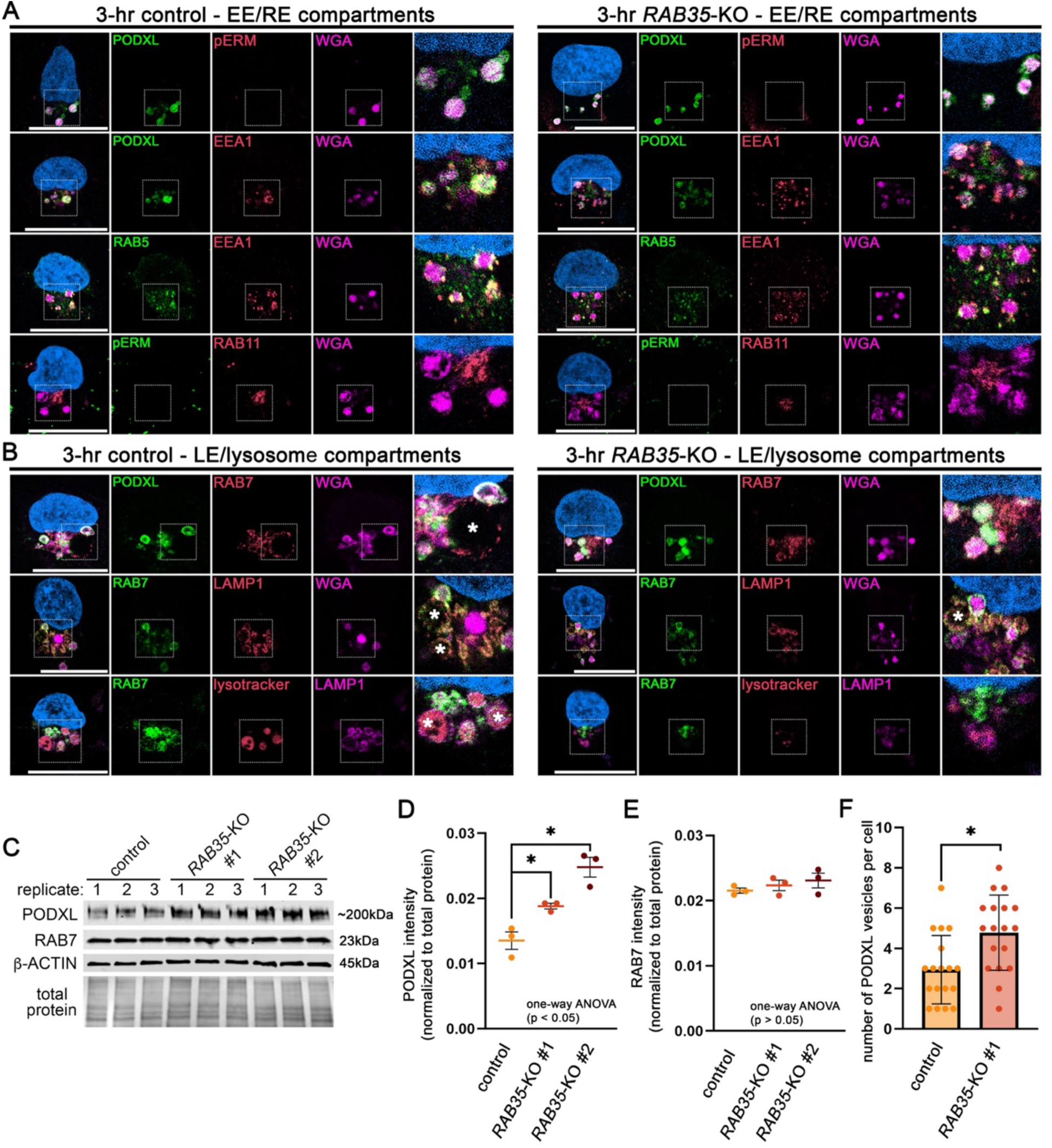
Loss of RAB35 impairs LE/lysosome formation during apicosome formation. A,B) Representative confocal images of 3-hr control and *RAB35*-KO cells stained with indicated markers (EE and RE compartments (A); LE/lysosome compartments (B)). Asterisks in (B) indicate highly enlarged LE/lysosomes that are labeled by RAB7, LAMP1 and/or lysotracker. C) Western blot analysis of 3-hr control, *RAB35*-KO #1 and *RAB35*-KO #2 cells, blotted for PODXL, RAB7 and β-actin. Three independent replicates are shown for each sample group. D,E) Quantitations for normalized levels of PODXL (D) and RAB7 (E), normalized against total protein, using samples in (C). One-way ANOVA analysis showed a statistical significance in D (PODXL, p < 0.05), but not in E (RAB7, p > 0.05); asterisks indicate a statistical significance based on Tuckey’s test (p < 0.05). F) Quantitation for the number of PODXL vesicles (widest diameter larger than 2µm) in 3-hr control and *RAB35*-KO cells (n = 18 cells per background, asterisk indicates a statistically significant difference based on Student’s t-test (p < 0.05)). Insets indicate magnified regions shown as merged images. Scale = 20µm. Blue pseudocolor indicates DNA in all images.

Moreover, the number of PODXL^+^ vesicles per cell were increased in the *RAB35*-KO cells at 3-hr (**Fig. 6F**). Together, these results reveal that the multiple apicosome formation in the *RAB35*-KO cells is associated with both defective LE/lysosome dynamics at the pre-apicosome stage and increased PODXL levels. Thus, RAB35 may function to drive lysosome formation, and/or to traffic PODXL to the lysosome.

We next explored the effects of the loss of RAB35 on the endosomes and lysosomes at 20-hr (**Fig. S6C,D**). At this timepoint, components of the EE, RE, LE and lysosomes were adjacent to the multiple PODXL and pERM positive apicosome-like structures in the absence of RAB35 (**Fig. S6C,D**). Therefore, although the *RAB35*-KO cells are unable to form a single mature apicosome structure, the pERM-positive apicosome-like structures do not display endosomal characteristics.

To test whether RAB35 GTPase activity contributes to these functions, we examined apicosome formation in *RAB35*-KO cells that inducibly express wild-type (WT), constitutively active (CA – Q67L), or dominant negative (DN – S22N) forms of GFP-RAB35 fusion protein. We found that expression of either RAB35-WT or -CA constructs restored single apicosome formation, while expression of RAB35-DN did not restore single apicosome formation (**Fig. 7A**, quantitation shown in **Fig. 7B,S7A**).

**Figure 7.**
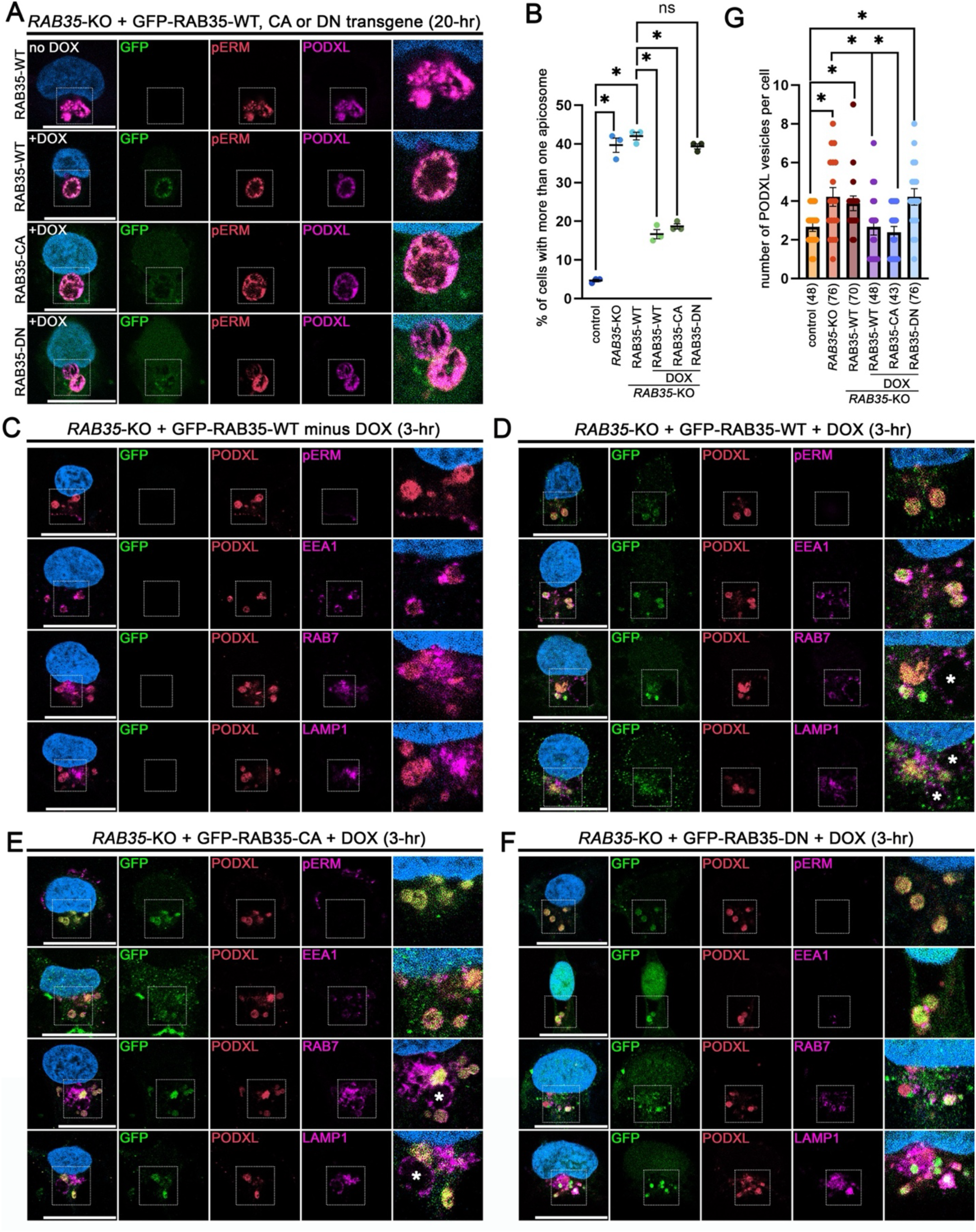
RAB35 GTPase activity is required for the formation of single apicosomes and lysosomal compartments. A) Representative confocal images of *RAB35*-KO cells carrying DOX-inducible GFP fused RAB35-WT,-CA (constitutively active, Q67L) and -DN (dominant negative, S22N) transgenic constructs treated with DOX for 20 hours, stained with pERM and PODXL. In the “no DOX” condition (top), DOX-untreated 20-hr *RAB35*-KO + RAB35-WT cells were harvested at the 20-hr timepoint. Note that the expression of the RAB35-DN construct does not lead to single apicosome formation. B) Quantitations for 20-hr cells with more than one apicosome for indicated background (n = 100 cells in each of the three independent experiments in each background. A statistical significance of the sample group was seen based on one-way ANOVA (p < 0.05) as well as by Tuckey’s test (asterisks indicate statistical significance of selected pairwise comparisons, p < 0.05)). C-F) Representative optical section of *RAB35*-KO cells carrying inducible GFP-RAB35-WT (DOX-untreated in (C); DOX-treated in (D)), -CA (E) and -DN (F) constructs, harvested at the 3-hr timepoint (with 3-hr DOX treatment) and stained with indicated markers. Note that, while seen with the expression of RAB35-WT or -CA, the expression of the RAB35-DN construct does not lead to the formation of highly enlarged RAB7/LAMP1 double positive LE/lysosomes. Asterisks indicate the highly enlarged LE/lysosome compartments (D,E). G) Quantitation for the number of PODXL vesicles (widest diameter larger than 2µm) in 3-hr cells from indicated sample group (n = 18 cells per background; numbers indicate total counted vesicles). Number of the PODXL vesicles is reduced to levels similar to controls when the RAB35-WT or -CA construct is expressed in the *RAB35*-KO background, which is not seen with the expression of the RAB35-DN. A statistical significance of the sample group was seen based on one-way ANOVA (p < 0.05) as well as by Fisher’s LSD test; selected pairwise comparisons are shown (asterisks, p < 0.05; ns, p > 0.05). Insets indicate magnified regions in the merged images. Scale = 20µm. Blue pseudocolor indicates DNA in all images.

We also examined the impact of RAB35-CA and -DN expression on endosome dynamics. When imaged at 3-hr, *RAB35*-KO cells expressing either RAB35-WT or RAB35-CA showed highly enlarged structures that were labeled with LE/lysosome components but lacked PODXL/WGA, similar to controls (**Fig. 7C-E**, see asterisks in **Fig. 7D**,**E**). In contrast, these large compartments were lacking in cells expressing RAB35-DN (**Fig. 7F**). Moreover, while RAB35-WT or -CA expression restored control-like PODXL vesicle formation, this rescue was not seen with RAB35-DN expression (**Fig. 7G**). Together, these results suggest that the GTPase activity of RAB35 is required for the formation of a single apicosome, and for the formation of the highly enlarged LE/lysosome compartment at the pre-apicosome stage.

### RAB35 acts upstream of RAB7 to promote single apicosome formation

Rab35 functions upstream of Rab5 and Rab7 in a phagosomal pathway in C. elegans (31, 32). Moreover, previous studies showed that Rab7 activity promotes LE trafficking and lysosome formation (33–36). Given the observation that the loss of RAB35 impairs the formation of the highly enlarged LE/lysosomes, we explored whether RAB35 acts upstream of RAB7 during apicosome formation. To test this hypothesis, we inducibly expressed WT, constitutively active (CA – Q67L), or dominant negative (DN – T22N) forms of GFP-RAB7 fusion protein in the *RAB35*-KO background. Strikingly, expression of GFP-RAB7-WT or -CA (but not DN) in *RAB35*-KO restored the formation of single apicosomes (**Fig. 8A**, quantitation in **Fig. 8B**,**S7B**), highly enlarged LE/lysosomes (**Fig. 8C-F**, asterisks), as well as PODXL vesicles (**Fig. 8G**). This result suggests that the RAB35-RAB7 axis controls single apicosome formation.

**Figure 8.**
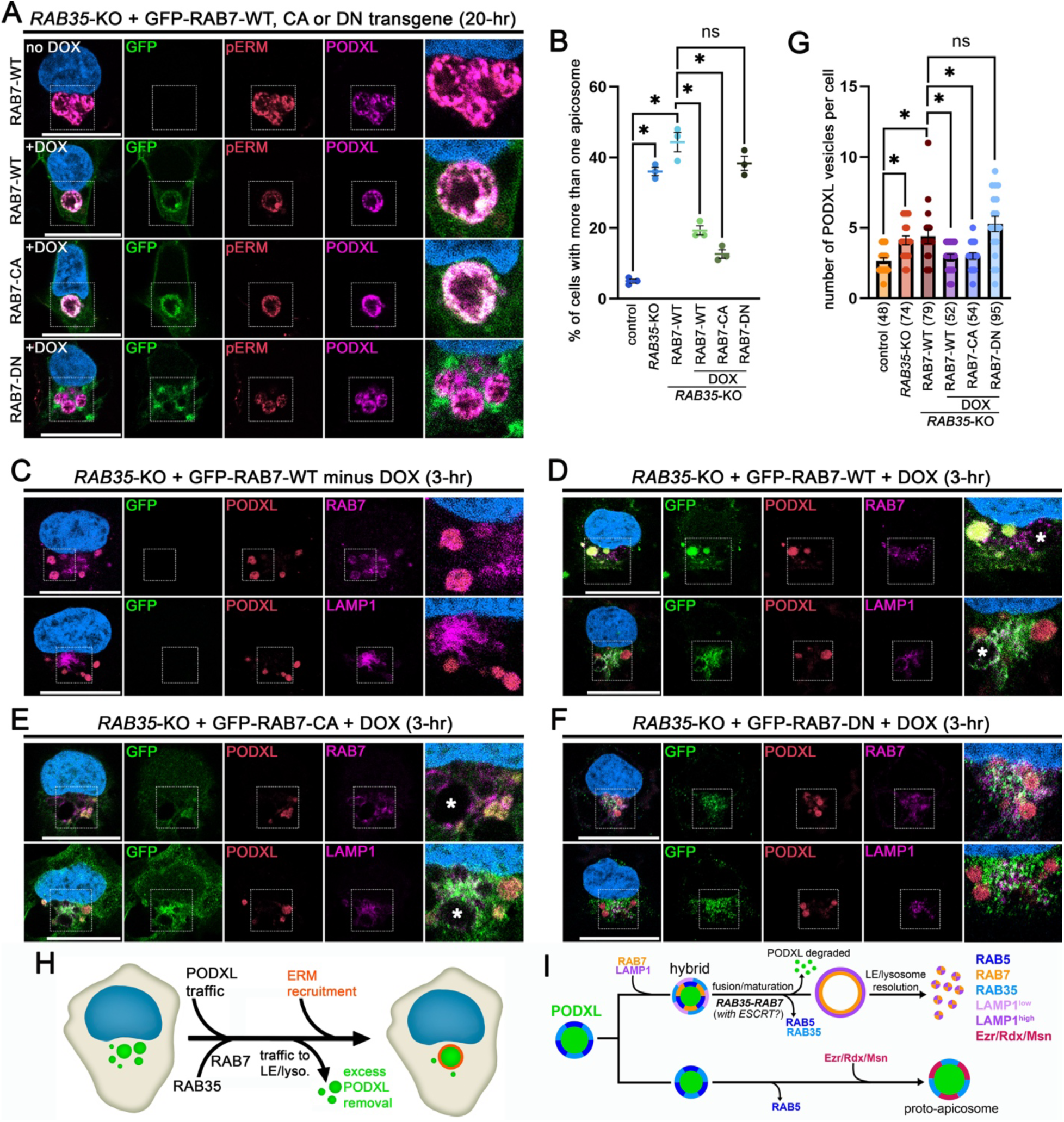
RAB7 acts downstream of RAB35 in apicosome formation and lysosome expansion. A) Representative optical section of *RAB35*-KO cells inducibly expressing GFP-RAB7-WT, -CA (Q67L) or -DN (S22N) construct at 20-hr (with 20-hr DOX treatment), stained with indicated markers. In the “no DOX” condition (top), DOX-untreated 20-hr *RAB35*-KO + RAB7-WT cells were harvested at the 20-hr timepoint. RAB7-WT or -CA expression leads to single apicosome formation while several apicosomes are still seen with RAB7-DN expression. B) Quantitation for the percentage of cells with more than one apicosome in the indicated genetic background (n = 100 cells in each of the three independent experiments in each background). Increase in RAB7 level or activity leads to reduced multi-apicosome formation. A statistical significance of the sample group was seen based on one-way ANOVA (p < 0.05) as well as by Tuckey’s test; selected pairwise comparisons are shown (asterisks, p < 0.05; ns, p > 0.05). C-F) Representative confocal images of *RAB35*-KO cells inducibly expressing GFP-RAB7-WT (minus DOX in (C), plus DOX in (D)), -CA (Q67L, (E)) or -DN (T22N, (F)) construct at 3-hr (with or without 3-hr DOX treatment), stained with indicated markers. Formation of then highly enlarged LE/lysosomal compartments (asterisks) is seen with RAB7-WT or -CA expression (D,E). G) Quantitation for the number of PODXL vesicles (widest diameter larger than 2µm) in 3-hr cells from indicated sample group (n = 18 cells per background; numbers indicate total counted vesicles). Number of the PODXL vesicles is reduced to levels similar to controls when the RAB7-WT or -CA construct (but not -DN) is expressed in the *RAB35*-KO background. A statistical significance of the sample group was seen based on one-way ANOVA (p < 0.05) as well as by Fisher’s LSD test; selected pairwise comparisons are shown (asterisks, p < 0.05; ns, p > 0.05). H,I) Proposed model of apicosome formation at whole cell (H) and vesicle (I) levels. RAB35-RAB7 machinery promotes removal of excess PODXL vesicles, and contributes to single apicosome formation (H). At the earliest stages of apicosome formation (I), an EE/LE hybrid vesicle population is formed from a group of PODXL vesicles labeled with RAB35 and RAB5 by recruiting RAB7 and LAMP1. Subsequently, these hybrid vesicles shed RAB35 and RAB5, and are trafficked to an acidic LE/lysosomal compartment in a RAB35-RAB7-dependent manner, where PODXL is degraded. Much of the highly enlarged LE/lysosome compartments are resolved by the time the proto-apicosome is formed (top). Another group of RAB35/RAB5 PODXL vesicles removes RAB5 and recruits ERM proteins, which contribute to the formation of the proto-apicosome (bottom). Insets indicate magnified regions in the merged images. * = statistically significant; ns = statistically insignificant. Scale = 20µm. Blue pseudocolor indicates DNA in all images.

## DISCUSSION

In this study, we examined endo-lysosomal dynamics during apicosome formation in a human epiblast model. We identified a hybrid compartment with early and late endosome-like characteristics, as well as previously unrecognized dynamics of late endosome and lysosome compartments in trafficking PODXL. We also demonstrated that RAB35 is a marker of the apicosome, and that RAB35 is critical for driving single apicosome formation as well as LE/lysosome dynamics. These findings provide key insights into how LE and lysosomal compartments contribute to human epiblast lumen formation and expand the list of machinery required for apicosome formation (**Fig. 8H**, (1, 8, 15, 37)).

Our results are consistent with a model whereby the apicosome derives from one or more of the enlarged RAB5 endosomes observed at early timepoints. At the earliest stages of apicosome formation, nearly all PODXL is associated with RAB5. This finding is consistent with our prior work, which suggests that the apicosome contains endocytosed materials (7). Since RAB5 endosome often serves as the first way-station for endocytosed cargo (23), it seems clear that the co-localization with RAB5 reflects the transit of PODXL through the early endosome. The enlarged nature of a subset of the RAB5 endosomes remains notable; however, whether this reflects their formation via a process like micropinocytosis, fusion of multiple smaller endosomes, or enlargement of individual endosomes via other processes remains unknown.

The hybrid enlarged RAB5-RAB7 organelle is a striking aspect of apicosome formation. At early stages of apicosome formation, many of the enlarged vesicles that contain PODXL and RAB5 also contain RAB7. RAB7 generally defines the LE and is often recruited by machinery that is recruited by RAB5 (38). Therefore, we postulate that a subset of the enlarged RAB5 endosomes recruit RAB7, and that these are destined for destruction, whereas those that do not recruit RAB7 ultimately contribute to the apicosome (**Fig. 8I**).

It remains unclear how the RAB5-RAB7 organelle forms and how it resolves. In many systems, RAB5 endosomes convert to a RAB7 endosome (39). During this conversion, there is a transient period of overlap (40). Based on the presence of abundant RAB5-RAB7 organelles observed in the asynchronous system described here using hPSC, it is difficult to envision that this is a transient event during apicosome formation. Therefore, it remains unclear whether the conversion from RAB5 to RAB7 is mechanistically different during hPSC-epiblast formation.

Our data argue that RAB35 may contribute to this conversion, because overexpression of RAB7 can suppress many of the effects associated with the loss of RAB35. Curiously, we note that ESCRT proteins have been recently linked to the RAB5-RAB7 conversion (41, 42). We find the ESCRT-0 proteins HGS (also known as HRS), STAM1 and STAM2 enriched in the PODXL-proximity proteome, suggesting that this aspect of RAB5-RAB7 conversion may be functional in hPSC during apicosome formation (**Fig. 8I**). Moreover, HGS has previously been shown to bind to RAB35, where it promotes degradation of synaptic vesicle proteins (43). Therefore, it is possible that RAB35 provides a linchpin for the RAB5-RAB7 conversion in hPSC via ESCRT-0.

Apicosome formation is only the first step of lumen formation in the hPSC-epiblast model. After formation in single cells, apicosomes in an aggregate of cells relocate to the center of the aggregate to initiate the formation of a central lumen in a cyst of cells (7, 44). Therefore, it will be of future interest to examine whether the multiple apicosome phenotype associated with the loss of RAB35 impacts the subsequent step of central lumen formation, as well as to investigate whether LE and lysosomal functions also contribute to this apicosome-driven lumenogenesis event. Taken together, this study opens an avenue to explore the role of LE and lysosome compartments during apical membrane morphogenesis.

## MATERIALS AND METHODS

### Cell lines used in this study

H9 hESC line was used in this study (WA09, P30, WiCell; National Institute of Health (NIH) registration number: 0062). All protocols for the use of the hESC line were approved by the Human Stem Cell Research Oversight Committee at the Medical College of Wisconsin. H9 hESC were maintained in a feeder-free system for at least 20 passages and characterized as karyotypically normal at several passage number including P30, P46 and P54. Karyotype analysis was performed at Cell Line Genetics. All hPSC lines tested negative for mycoplasma contamination (LookOut Mycoplasma PCR Detection Kit, Sigma-Aldrich). All transgenic and KO hPSC lines in this study used H9 hESC as the parental line.

hESC were maintained in a feeder-free culture system with 50%/50% mix of mTeSR1 and mTeSR plus (STEMCELL Technologies). H9 cells were cultured on 1% (v/v) Geltrex (Thermo Fisher Scientific), or with Cultrex SCQ (Bio-Techne) coated six-well plates (Nunc). Cells were passaged as small clumps every 4 to 5 days with Dispase (Gibco). All cells were cultured at 37°C with 5%CO_2_. Media was changed every day. hESC were visually checked every day to ensure the absence of spontaneously differentiated mesenchymal-like cells in culture. Minor differentiated cells were scratched off the plate under a dissecting scope once identified. The quality of all hESC lines was periodically examined by immunostaining for pluripotency markers and successful differentiation to three germ layer cells.

### Apicosome formation assays and staging

Methods for these assays are previously as described (7, 27, 45). Briefly, singly dissociated cells were prepared using Accutase (Sigma-Aldrich) and were plated on coverslips coated with 1% Geltrex at 1 x 10^4^ cells/cm^2^. Cells were plated in the 50/50 mTeSR1/mTeSR plus media containing Y-27632 (STEMCELL Technologies) and 2% Geltrex. Apicosome formation initiates spontaneously after plating.

Pre-, proto- and mature apicosome stages were identified based on the number, size, morphology and molecular characteristics (e.g., pERM labeling) of PODXL/WGA vesicles. In the pre-apicosome cells, three to four PODXL vesicles that are between 2-2.5µm in diameter at the widest point are seen. These PODXL vesicles lack labeling by pERM. In the proto-apicosome cells, in addition to one to two PODXL vesicles that are 2-2.5µm in diameter, one larger PODXL vesicle (3-3.4µm in diameter) that is labeled with pERM is present, which is the proto-apicosome. Mature apicosomes (positive for both PODXL and pERM) are larger (ranging between 6.7-8µm in diameter) and show a central open domain.

### Immunostaining

Samples were fixed using 4% paraformaldehyde at indicated timepoints for 40 to 60 min, then were rinsed with PBS three times, and permeabilized with 0.1% SDS (Sigma-Aldrich) solution for 40 min. The samples were blocked in 4% heat-inactivated goat or donkey serum (Gibco) in PBS for 1 hour to overnight at 4°C. The samples were incubated with primary antibody solution prepared in blocking solution at 4°C overnight, washed three times with PBS (10 min each), and incubated in blocking solution with goat or donkey raised Alexa Fluor–conjugated secondary antibodies (Thermo) at room temperature for 2 hours. Counterstaining was performed using Hoechst 33342 (nucleus, Thermo) or Alexa Fluor–conjugated WGA (membrane, Thermo). For LysoTracker (LysoTracker Red DND-99, 500nM in DMSO) staining, cells were treated with LysoTracker for 30 minutes before harvesting, and were processed for immunostaining. All samples were mounted on slides using Fluoromount-G (Thermo). Antibodies for IF staining are found in **Table 3**.

**Table 3.** List of reagents used in this study. (A-D) List of antibodies (A), primers (B), obtained plasmid (C) and newly generated plasmid (D) used in this study.

### Confocal microscopy

Confocal images were acquired using a Zeiss LSM980 laser scanning confocal microscope. Zen (Zeiss) as well as Photoshop (Adobe) was used to process images.

### Constructs and cell lines

Generation of the PODXL-APEX2 and APEX2-NES constructs has been previously described (15). piggyBac-based DOX inducible RAB35 (Addgene#47424, 47425, 47426 (gift of Peter McPherson (46))) and RAB7 (Addgene#12605 (gift of Richard Pagano (47)), 28048, 28049 (gift of Qing Zhong (48))) constructs were generated by PCR amplification (forward primer: Clo-dTOPO-EGFP-fw; reverse primers: Clo_dTOPO_hRab35_rv (RAB35), Clo_dTOPO_EGFP-RAB7A-rv (RAB7)) and by subcloning the PCR products into the pENTR-dTOPO (Thermo), followed by Gateway cloning (Thermo) into the PB-TA-ERN (Addgene#80474, gift of Knut Woltjen (49)).

### piggyBac-based transgenic and genome-edited hESC lines

The pBACON-puro-h*RAB35* construct used to generate an additional *RAB35*-KO line has been established previously (15). To generate transgenic or genome-edited hESC lines, piggyBac constructs (4µg) and pCAG-ePBase (1µg; gift from Ali Brivanlou) were cotransfected into H9 hESC (60,000 cells/cm^2^) using GeneJammer transfection reagent (Agilent Technologies). To enrich for cells expressing the construct, drug selection (puromycin, 2 µg/ml; neomycin, 250 µg/ml) was performed 48 to 72 hours after transfection. Selected pools were plated sparsely, and small colonies were picked for further expansion to establish clonal lines. hESC stably expressing each construct maintained the expression of pluripotency markers.

During pBACON-based genome editing, puro-selected cells were cultured at low density (300 cells/cm^2^) for clonal selection. Established colonies were manually picked and expanded for screening indel mutations using PCR amplification of a region spanning the targeted gRNA region (primer pair: Seq_hRAB35_fw and Seq_hRAB35_rv), which were subcloned into pPBCAG-GFP (gift of Joseph Loturco, (50)) at Eco RI and Not I sites, and sequenced (Seq-3′ TR-pPB-Fw). Genomic DNA was isolated from individual clones using DirectPCR Lysis Reagent (Tail) (VIAGEN). At least 12 to 15 bacterial colonies were sequenced to confirm genotypic clonality. Control cells are H9 hESC in all loss-of-function experiments.

### APEX2 labeling and sample preparation

To examine APEX2 labeling, singly dissociated PODXL-or NES-APEX2 cells were plated on glass coverslips at 1 x 10^4^ cells/cm^2^ in mTeSR1/mTeSR plus media containing DOX (2 µg/ml) and Geltrex (2%) for 29 hours. In the last 1 hour, cells were incubated in mTeSR/mTeSR plus media containing 2% Geltrex and biotin-phenol (BP, biotinyl tyramide, 500µM, AdipoGen) without DOX. Hydrogen peroxide (H_2_O_2_) was then added directly into the medium to a final concentration of 1mM for 90 seconds at room temperature to initiate biotinylation, and cells were immediately fixed using 4% paraformaldehyde for fluorophore conjugated streptavidin staining as well as immunostaining for microscopic analysis.

To prepare proteomics samples, the 6 samples were prepared individually (shown in **Fig. 3A**) in 10 tissue culture-treated 100mm dishes (thermo, precoated with 1% Geltrex). For each plate 1.0 x 10^6^ hPSC (DOX-inducible stable lines) were plated to obtain approximately 1.0 to 1.2 x 10^7^ cells per sample at 30-hr, sufficient for 3.0 mg of total protein. At 30-hr, APEX2 labeling was performed as described above. In addition, followed by the H_2_O_2_ treatment step, cells were washed 3x using quencher solution (10mM sodium ascorbate (Spectrum Chemical), 10mM sodium azide (Sigma-Aldrich) and 5mM Trolox (Sigma-Aldrich) in Dulbecco’s PBS (Gibco)).

After quenching, APEX2-labeled samples from 10 dishes were then resuspended as a pool in quencher buffer and centrifuged at 500g for 5 minutes to collect the cell pellet for lysis in radioimmunoprecipitation assay (RIPA) lysis buffer (Pierce) containing 1x Halt protease inhibitor cocktail (Thermo), 1mM phenylmethylsulfonyl fluoride (Sigma-Aldrich), 10mM sodium azide, 10mM sodium ascorbate and 5mM Trolox. Cell lysates were centrifuged at 15,000g for 15min at 4°C, and supernatant was collected for enriching biotinylated proteins using streptavidin beads.

The Pierce 660-nm assay (Pierce) was used to quantify protein concentrations in sample supernatants. To isolate biotinylated proteins, streptavidin-coated magnetic beads (Pierce) were first washed twice with RIPA lysis buffer. For each sample, 3.0 mg of total protein was incubated with 500 µl of streptavidin beads overnight at 4°C with gentle rotation. Beads were subsequently washed on a MagnaRack (Thermo Fisher Scientific). Beads were washed twice with RIPA lysis buffer, once with 1 M KCl, once with 0.1 M Na_2_CO_3_, once with 2 M urea in 10 mM tris-HCl (pH 8.0), then twice with RIPA lysis buffer, and three times with PBS. Last, PBS was removed as much as possible and beads were frozen in −80°C before performing on-bead digestion, TMT labeling, peptide pooling, fractionation, and LC-MS/MS. All these steps were performed at 4°C unless otherwise noted.

### Western blot

SDS-PAGE gels (10% or gradient gel, 4 to 20%, Bio-Rad) and polyvinylidene difluoride membranes were used. Membranes were blocked using Intercept (TBS) Blocking Buffer (LI COR), total protein quantification was performed by using Revert 700 Total Protein Stain (LI COR), and primary antibody overnight incubation was performed at 4°C, followed by 2-hour IRDye (LI-COR) secondary antibody incubation. Biotinylated proteins were detected and quantified by streptavidin-IRDye conjugate (LI-COR). Blots were imaged using LI-COR Odyssey Infrared Imaging system.

### Quantitative MS

Proteins bound to streptavidin beads were digested by trypsin following the standard on-bead trypsin digestion workflow (51). Samples were proteolyzed and labeled with TMTsixplex by following the manufacturer’s protocol (Thermo Fisher Scientific) with minor modifications. Briefly, upon reduction [10 mM DTT in 0.1 M Triethylammonium bicarbonate (TEAB); 45°C, 30 min] and alkylation (55 mM 2-chloroacetamide in 0.1 M TEAB; room temperature, 30 min in dark) of cysteines, the proteins were digested overnight with trypsin (1:25; enzyme:protein) at 37°C, with constant mixing using a thermomixer. Proteolysis was stopped by adding 0.2% trifluoroacetic acid and peptides were desalted using SepPak C18 cartridge (Waters Corp). The desalted peptides were dried in Vacufuge (Eppendorf) and reconstituted in 100µl of 0.1 M TEAB. The TMTsixplex reagents were dissolved in 41µl of anhydrous acetonitrile, and labeling was performed by transferring the entire digest to TMT reagent vial and incubating at room temperature for 1 hour. The reaction was quenched by adding 8 µl of 5% hydroxyl amine and further 15-min incubation. Labeled samples were mixed together and dried using a vacufuge. An offline fractionation of the combined sample into six fractions was performed using a high-pH reversed-phase peptide fractionation kit according to the manufacturer’s protocol (Pierce; catalog no. 8488). Fractions were dried and reconstituted in 12 µl of 0.1% formic acid/2% acetonitrile in preparation for LC-MS/MS analysis.

To improve quantitation accuracy, we used multinotch-MS3 (51), which minimizes the reporter ion ratio distortion resulting from fragmentation of co-isolated peptides during MS analysis. Orbitrap Fusion (Thermo Fisher Scientific) and RSLC Ultimate 3000 nano-UPLC (Dionex) were used to acquire the data. The sample (2 µl) was resolved on a PepMap RSLC C18 column (75 µm inside diameter × 50 cm; Thermo Fisher Scientific) at a flow rate of 300 nl/min using 0.1% formic acid/acetonitrile gradient system (2 to 22% acetonitrile in 110 min; 22 to 40% acetonitrile in 25 min; 6-min wash at 90% followed by 25-min re-equilibration) and directly sprayed onto the mass spectrometer using EasySpray source (Thermo Fisher Scientific). The mass spectrometer was set to collect one MS1 scan (Orbitrap; 120,000 resolution; AGC target, 2 × 10^5^; max IT, 50 ms) followed by data-dependent, “Top Speed” (3 s) MS2 scans (collision-induced dissociation; ion trap; NCD 35; AGC, 5 × 10^3^; max IT, 100 ms). For multinotch-MS3, top 10 precursors from each MS2 were fragmented by higher-energy-collisional-dissociation (HCD) followed by Orbitrap analysis [NCE 55; 60,000 resolution; AGC, 5 × 10^4^; max IT, 120 ms;100 to 500 *m/z* (mass/charge ratio) scan range].

### Ratiometric analysis of proteomic data

Proteome Discoverer (v2.1; Thermo Fisher Scientific) was used for initial data analyses. MS2 spectra were searched against the SwissProt human protein database (downloaded on 4 December 2018; 20331 reviewed entries) using the following search parameters: MS1 and MS2 tolerance were set to 10 parts per million and 0.6 Da, respectively; carbamidomethylation of cysteines (57.02146 Da) and TMTlabeling of lysine and N termini of peptides (229.16293 Da) were considered static modifications; oxidation of methionine (15.9949 Da) and deamidation of asparagine and glutamine (0.98401 Da) were considered variable. Identified proteins and peptides were filtered to retain only those that passed ≤1% FDR threshold and ≥2 unique peptides. Quantitation was performed using high-quality MS3 spectra using the Reporter Ion Quantifier Node of Proteome Discoverer (average signal-to-noise ratio of 10 and <30% isolation interference). Specific TMT ratios were normalized using known 495 mitochondria matrix soluble proteins (17): 265 were found in the PODXL territory dataset for ratiometric analyses in Filter 1 [PODXL-APEX2 #1/negative (126/130), PODXL-APEX2 #2/negative (127/130) and in Filter 2 [PODXL-APEX2 #1/APEX2-NES #1 (126/128), PODXL-APEX2 #2/APEX2-NES #2 (127/129). In each TMT ratio, the median ratio of 265 mitochondrial matrix soluble proteins was calculated (PODXL-APEX2 #1/negative, 3.09; PODXL-APEX2 #2/negative, 2.69; PODXL-APEX2 #1/APEX2-NES #1, 1.06; PODXL-APEX2 #2/ APEX2-NES #2, 0.93); all proteins in each TMT ratio were divided using these values to generate normalized PODXL territory dataset (ratios in log2 scale). In Filter 1 (F1) and Filter 2 (F2), true positive (TP-F1 and TP-F2, proteins with UniProt “cell membrane” and “plasma membrane” annotations) and false positive [FP, 265 mitochondrial matrix soluble proteins (Filter 1) and all proteins in the list lacking UniProt “cell membrane” and “plasma membrane” annotations (Filter 2)] were defined. TP-F1 and TP-F2 are proteins that are known to localize in the membrane territory; FP-F1 are proteins that are predicted to be nonbiotinylated by APEX2 fusion constructs in this study; FP-F2 consists of proteins that can be biotinylated by our APEX2 constructs but are not predicted to be proximal to the PODXL territory. True-positive rate (TPR) and false-positive rate (FPR) were calculated for each ratio [TPR = TP/(TP + FN); FPR = FP/(FP + TN); FN, false negative; TN, true negative]; these values were used to generate ROC curves to test the suitability of TP and FP for each ratiometric analysis based on the area under the curve (AUC; a commonly used statistic that calculates the area under the ROC curve and quantifies the probability in which a randomly chosen positive case outranks a randomly chosen negative case), as well as to determine cutoffs at which the largest difference between TPR and FPR was observed (PODXL-APEX2 #1/negative [log2(126/130) – AUC = 0.97, cutoff = 0.551]; PODXL-APEX2 #2/negative [log2(127/130) – AUC = 0.97, cutoff = 0.702]; PODXL-APEX2 #1/APEX2-NES #1[log2(126/128) – AUC = 0.7, cutoff = 0.19]; PODXL-APEX2 #2/APEX2-NES #2 [log2(127/129) – AUC = 0.74, cutoff = 0.062]) (**Fig. 3D**). TPR-FPR is equivalent to the Youden index (52), which is a statistic commonly used to represent the performance of a dichotomous test. Larger values of the index mean better performance. For example, a value of 1 would mean the performance is perfect as there are no false positives or false negatives. During the analysis of filter 2, proteins that passed filter 1 were used.

Given a threshold parameter *T*, we have AUC and cutoff (CO) formulas as

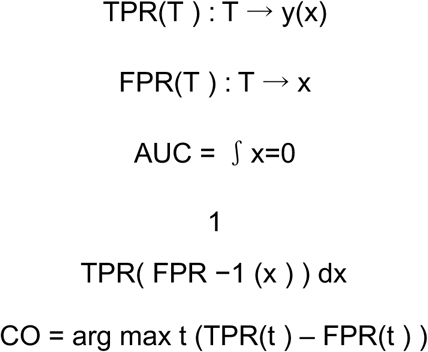

### GO enrichment analysis

The final apical (322) proteomes were uploaded to the STRING database [string-db.org (53), analysis performed on December 2024]: The top GO terms (ranked by FDR) on Cellular Component, Biological Process, and Molecular Function were plotted (**Fig. 3F-H**).

### Quantitative analyses

Graphs were generated using Prism 6 (GraphPad Software), and Student’s test (**Fig. 6F**,**S6B**) or one-way ANOVA (**Fig. 5B,6D,6E,7B,7G,8B,8G,S6A,S7**, followed by Tuckey or Fisher’s LSD post-hoc test where appropriate) was performed to examine statistical significance. p > 0.05 was considered nonsignificant; p ≤ 0.05 was considered significant and marked with an asterisk.

For **Fig. S1B**, 100 cells were counted from three independent samples (total of 300 cells) per timepoint. For **Fig. 5B**, **7B**, **8B**, **S6A** and **S7**, 100 cells were counted from three independent samples (total 300 cells) per experimental condition. In **Fig. 6F**, **7G** and **8G**, 18 cells were counted per experimental background. For **Fig. S6B**, PODXL negative LAMP1 vesicle diameter was measured from 20 cells per genetic background using Imaris (Oxford Instruments).

## ACKNOWLEDGEMENTS

We thank K. Woltjen (Kyoto University) for the piggyBac DOX-inducible vector (Addgene, #80474 (49)), P. McPherson (McGill University) for human RAB35 (Addgene#47424, #47425, #47256 (46)), R. Pagano for human RAB7 wild-type (Addgene #12605 (47)), Q. Zhong for human RAB7 mutants (Addgene #28049, #28048 (48)), and A. Brivanlou (Rockefeller University) for ePiggyBac transposase and DOX-inducible ePiggyBac constructs. We also thank L. Juga for her technical support throughout the project. Funding: This work was supported by NIH grants R01-HD098231, R01-HD102496, and R01-GM129255; MCW CBNA Start-up funds; MCW Cancer Center Graduate Fellowship. We thank the University of Michigan Proteomics Resource Facility.

## Author contributions

A.R., S.W., M.C.D., and K.T. designed experiments; A.R., S.W., C.-W.L., A.E.C., L.E.T., N.S. and K.T. performed experiments; A.R., S.W., M.C.D., and K.T. analyzed data and wrote the manuscript; M.C.D., and K.T. supervised the project; all authors contributed to the manuscript.

## Data and materials availability

All data needed to evaluate the conclusions in the paper are present in the materials in the manuscript. Additional information and reagents can be provided by the authors pending scientific review and a completed material and transfer agreement, where appropriate: Requests should be directed to M.D. and K.T.

## Supplementary Figures and Figure Legends

**Figure S1.**
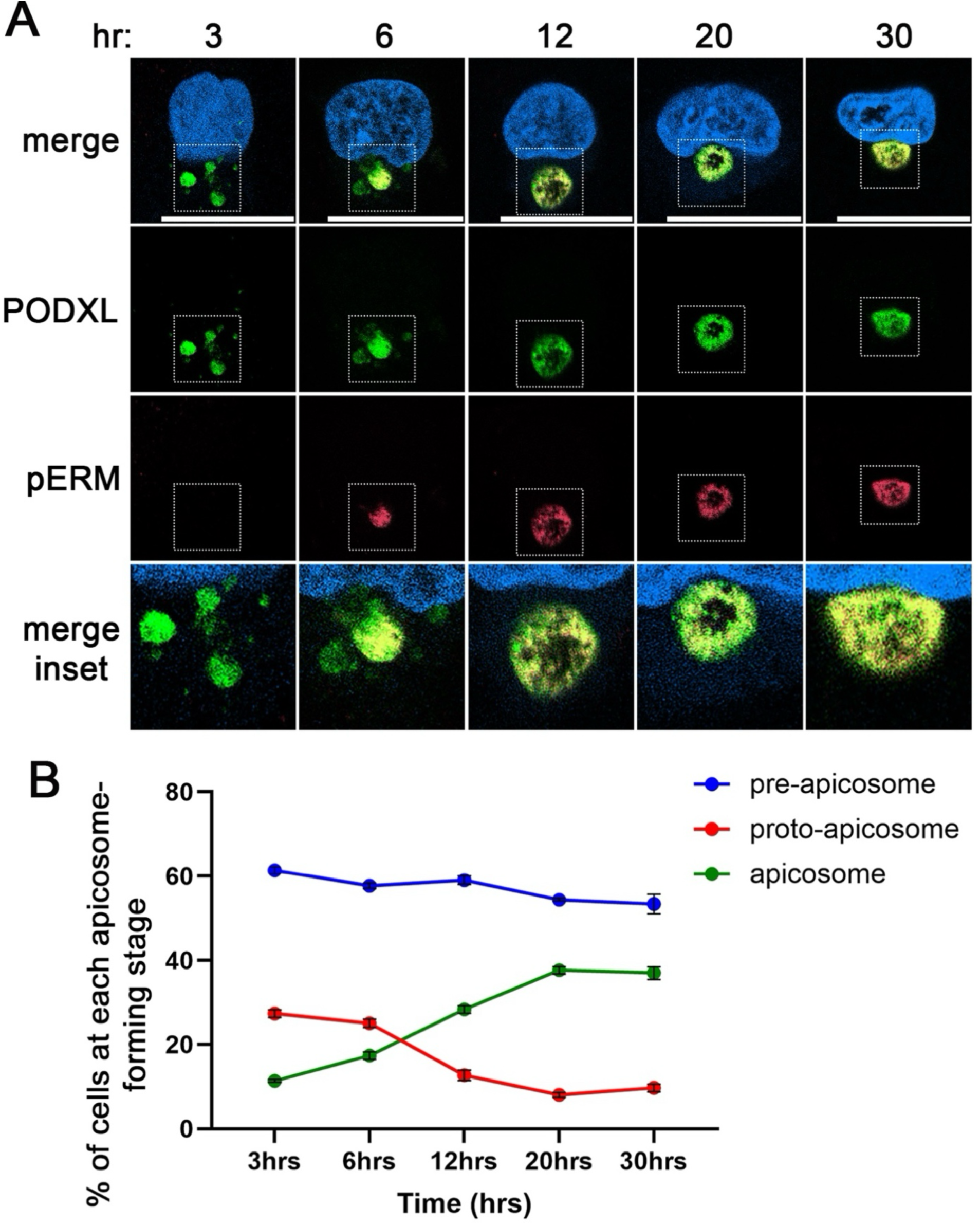
IF and quantitation analyses for apicosome formation at different timepoints. A) Representative optical sections of H9 hESC fixed at indicated timepoints, and stained for indicated markers. Insets that indicate magnified regions are included to better display trafficking dynamics and/or apicosome formation in the peri-nuclear domain. B) Quantitation for pre-apicosome, proto-apicosome and apicosome formation stages at indicated timepoints (n = 100 cells from each of the three independent experiments at each timepoint). Note that, over time, cells undergo cell division with or without apicosome formation; cells with the apicosome form a central lumen at the 2-cell stage. Since apicosome formation is reactivated in cells that have undergone mitosis without apicosomes, this accounts for the lack of reduction in pre-apicosome stage over time. Scale = 20µm. Blue pseudocolor indicates DNA in all images.

**Figure S2.**
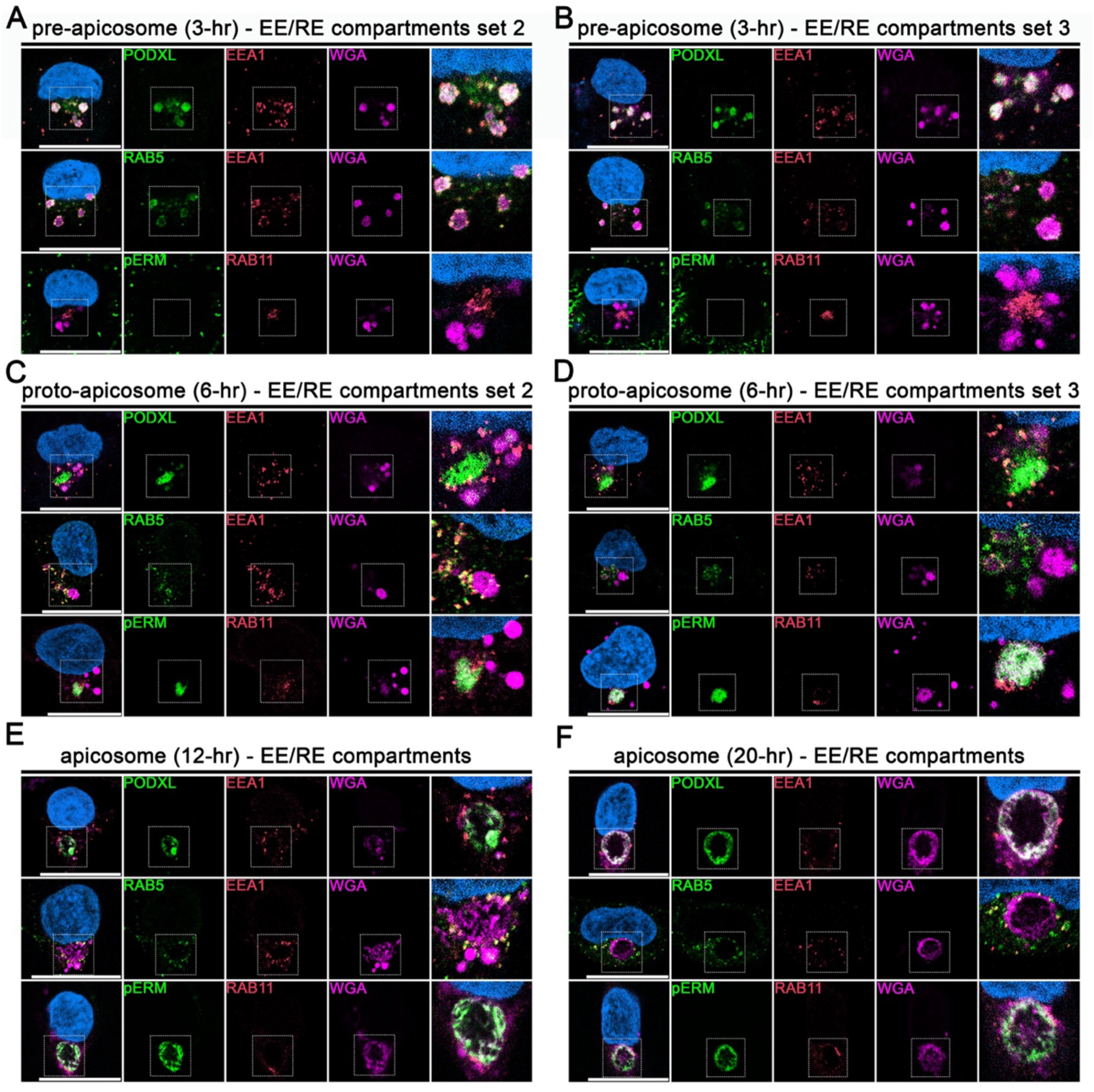
Additional IF images for EE and RE markers: second and third representative images for pre-(3-hr, A and B) and proto-(6-hr, C and D) apicosome cells in Fig. 1B and 1C, respectively, as well as representative images for cells with mature apicosome at 12-(E) and 20-(F) hr. Stained with indicated markers. Selected markers are shown. Insets indicate magnified regions in the merged images. Scale = 20µm. Blue pseudocolor indicates DNA in all images.

**Figure S3.**
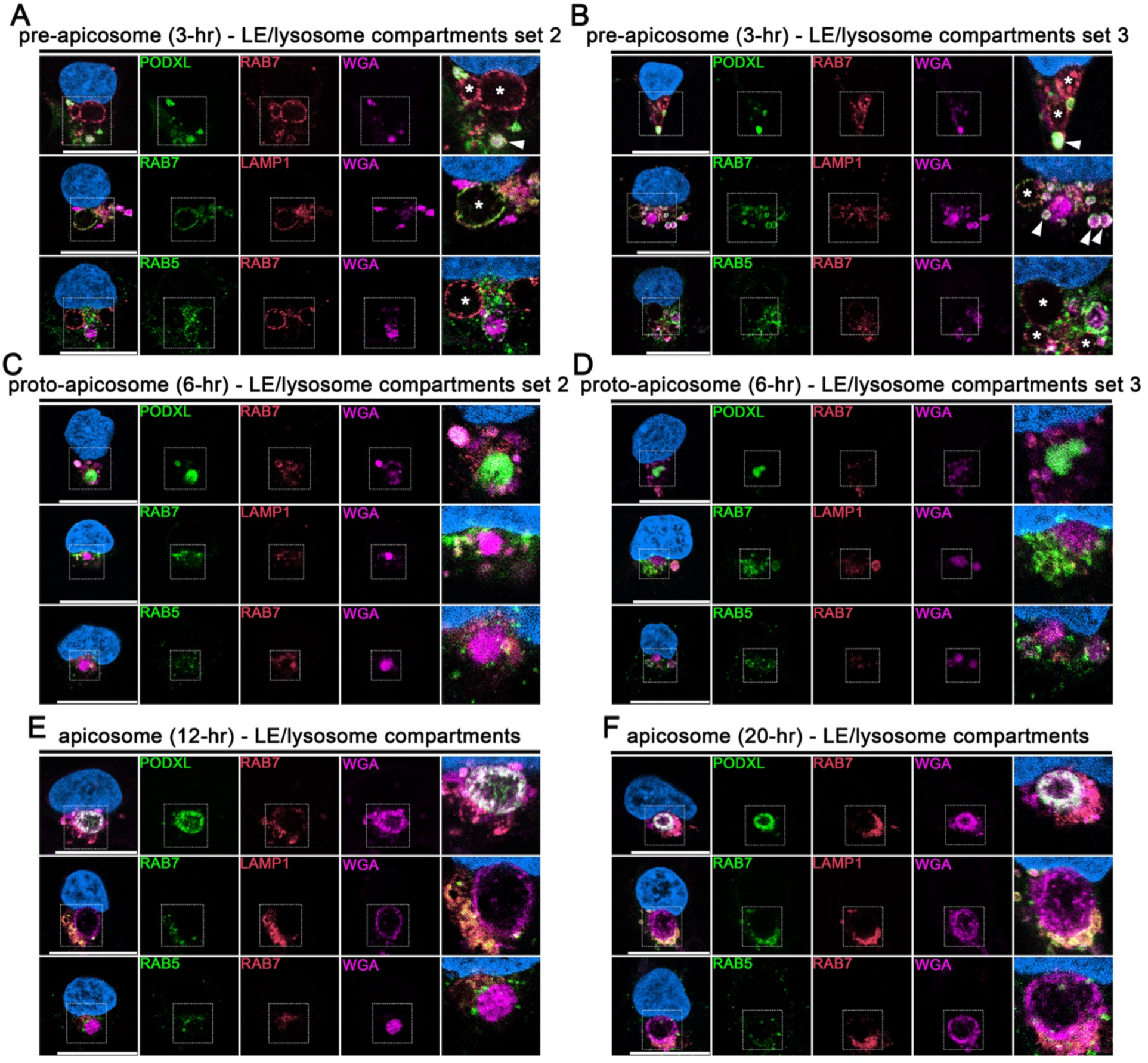
Additional IF images for LE/lysosome markers: second and third representative images for pre-(3-hr, A and B) and proto-(6-hr, C and D) apicosome cells in Fig. 1D and 1E, respectively, as well as representative images for cells with mature apicosome at 12-(E) and 20-(F) hr. Stained with indicated markers. Selected markers are shown. Insets indicate magnified regions in the merged images. Scale = 20µm. Blue pseudocolor indicates DNA in all images. Arrowheads indicate RAB7-labeled PODXL/WGA vesicles; asterisks indicate highly enlarged LE/lysosome compartments labeled by RAB7 and/or LAMP1. Scale = 20µm. Blue pseudocolor indicates DNA in all images.

**Figure S4.**
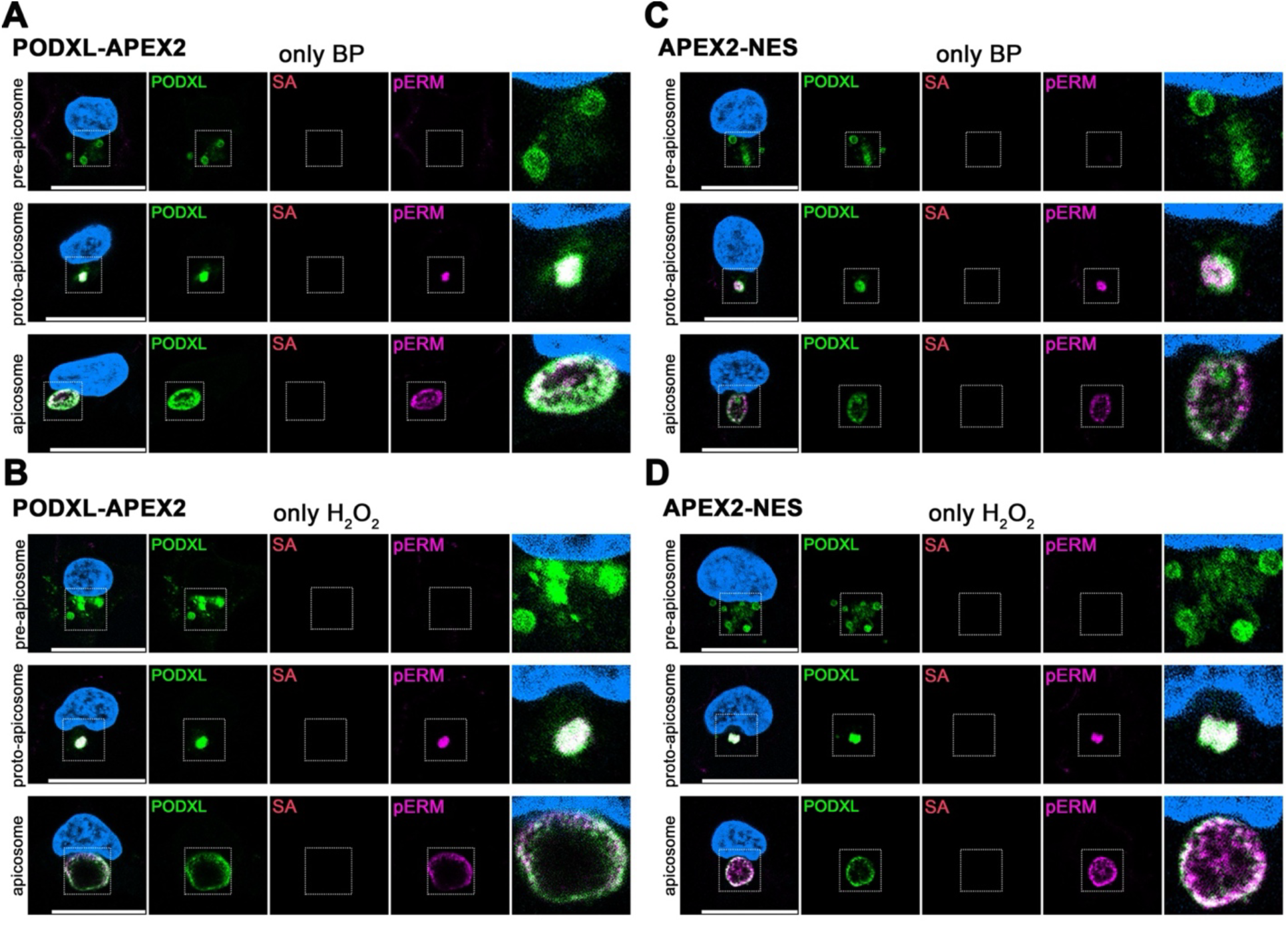
APEX2 labeling analysis of negative control samples. Representative optical sections of the DOX-treated PODXL-APEX2 (A,B) or APEX2-NES (C,D) cells in BP only (no H_2_O_2_, in A and C) or H_2_O_2_ only (no BP, in B and D) conditions from different apicosome forming stages, stained with indicated markers. Omitting either BP or H_2_O_2_ abolished the biotinylated signal. Insets indicate magnified regions in the merged images. Scale = 20µm. Blue pseudocolor indicates DNA in all images.

**Figure S5.**
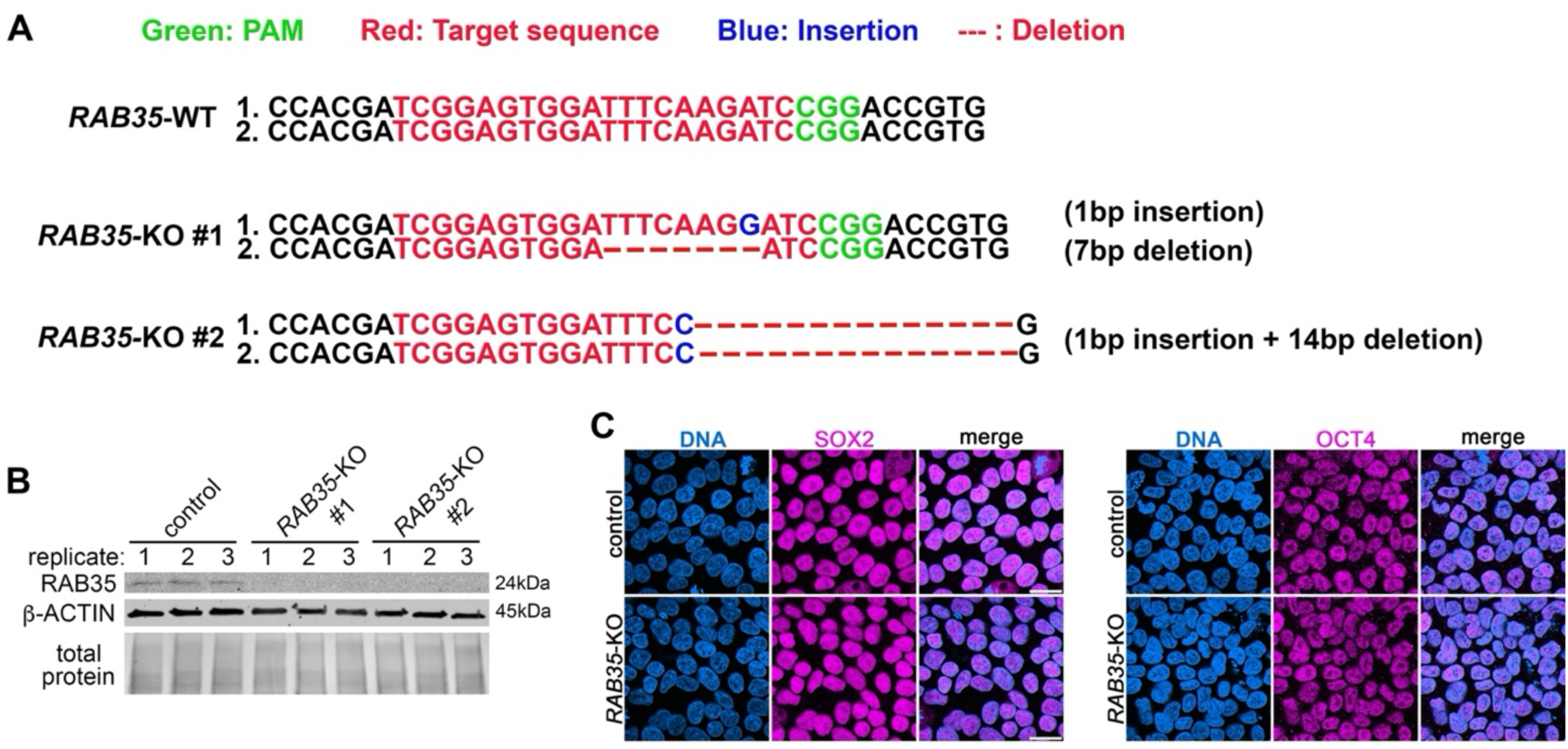
Design and validation of H9 hESC *RAB35* knockout lines. (A) Sequence of original (WT) and *RAB35*-KO lines. PAM sequence (green), target sequence (red), insertion (blue) and deletion (red dashed lines) are as indicated. (B) Western blot analysis for control, *RAB35*-KO #1 and *RAB35*-KO #2 lines. RAB35 protein is not detected in the two *RAB35*-KO lines. (C) Representative optical sections of control and *RAB35*-KO pluripotent monolayer samples, stained with SOX2 and OCT4, pluripotency markers. After 15 passages since genome editing, pluripotency is maintained. Scale = 20µm.

**Figure S6.**
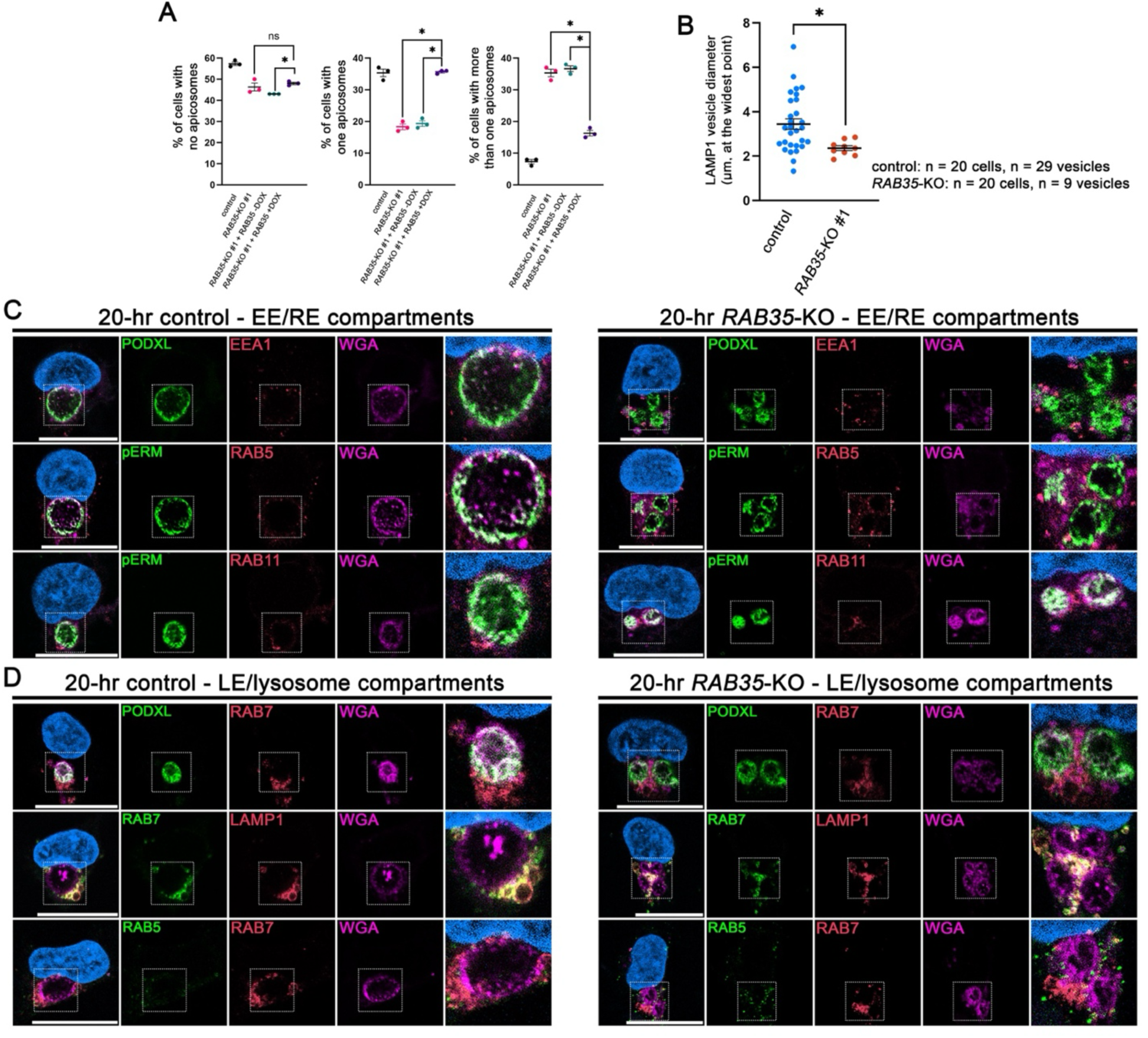
Additional characterization of *RAB35*-KO cells. (A) Quantitations for 20-hr cells with no apicosome (left, pre-apicosome stage), single apicosome (middle), or more than one apicosomes (right, same as the graph shown in Fig. 5B) in control (unmodified H9 hESC, black), *RAB35*-KO (red), and *RAB35*-KO + GFP-RAB35-WT without (green) or with (purple) DOX treatment (n = 100 cells in each of the three independent experiments in each background, statistical significance based on one-way ANOVA (p < 0.05) as well as by Tuckey’s test (asterisks indicate statistical significance of selected pairwise comparisons, p < 0.05)). (B) Quantitation for the diameter at the widest point of PODXL negative LAMP1^+^ LE/lysosome vesicles in control (blue) as well as in *RAB35*-KO (red) cells at 3-hr. Twenty pre-apicosome cells were counted for each background. Only nine PODXL negative LE/lysosome compartments were found in the KO background (among 20 total cells); twenty nine LE/lysosome compartments were found among 20 cells in the control background. Statistical significance (asterisk, p < 0.05) is based on Student’s t-test. (C,D) Representative confocal images of 20-hr control and *RAB35*-KO cells stained with selected apical (PODXL, WGA, pERM), EE (RAB5, EEA1), RE (RAB11), LE/lysosome (RAB7, LAMP1) markers. Insets indicate magnified regions in the merged images. Scale = 20µm. Blue pseudocolor indicates DNA in all images.

**Figure S7.**
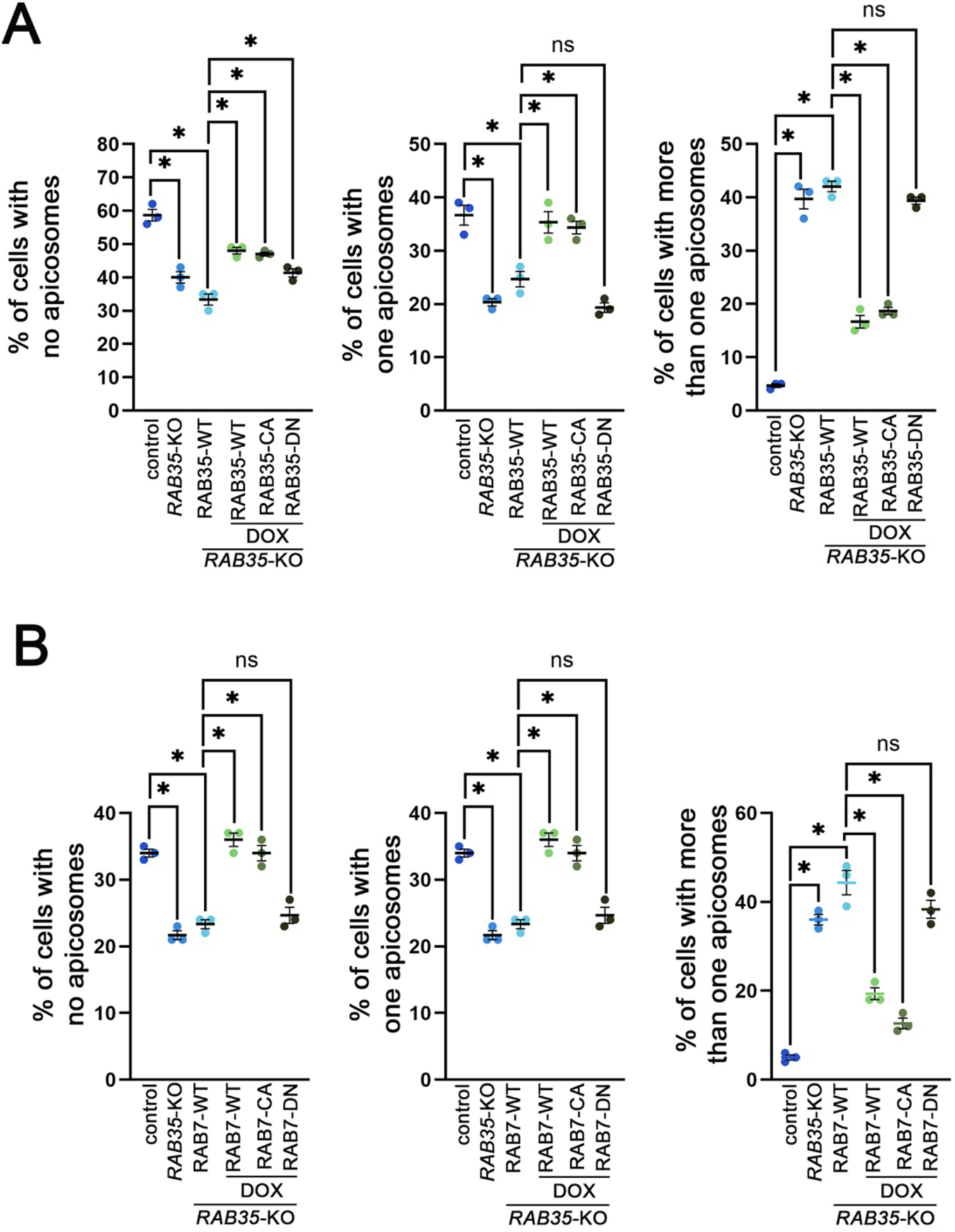
Apicosome formation quantitation analyses in Fig. 7B and 8B including all categories (no apicosome, one apicosome, more than one apicosome). (A) Quantitations for 20-hr cells with no apicosome (left, pre-apicosome stage), single apicosome (middle), or more than one apicosomes (right, same as the graph shown in Fig. 7B) in control (unmodified H9 hESC), *RAB35*-KO, *RAB35*-KO + GFP-RAB35-WT without DOX treatment, as well as DOX-treated *RAB35*-KO + GFP-RAB35-WT, *RAB35*-KO + GFP-RAB35-CA and *RAB35*-KO + GFP-RAB35-DN cells (n = 100 cells in each of the three independent experiments in each background). A statistical significance of the sample group was seen based on one-way ANOVA (p < 0.05) as well as by Fisher’s LSD test; selected pairwise comparisons are shown (asterisks, p < 0.05; ns, p > 0.05)). (B) Quantitations for 20-hr cells with no apicosome (left, pre-apicosome stage), single apicosome (middle), or more than one apicosomes (right, same as the graph shown in Fig. 8B) in control (unmodified H9 hESC), *RAB35*-KO, *RAB35*-KO + GFP-RAB7-WT without DOX treatment, as well as DOX-treated *RAB35*-KO + GFP-RAB7-WT, *RAB35*-KO + GFP-RAB7-CA and *RAB35*-KO + GFP-RAB7-DN cells (n = 100 cells in each of the three independent experiments in each background). A statistical significance of the sample group was seen based on one-way ANOVA (p < 0.05) as well as by Fisher’s LSD test; selected pairwise comparisons are shown (asterisks, p < 0.05; ns, p > 0.05)).

